# The Nesprin-1/-2 ortholog ANC-1 regulates organelle positioning in *C. elegans* without its KASH or actin-binding domains

**DOI:** 10.1101/2020.07.14.202838

**Authors:** Hongyan Hao, Shilpi Kalra, Laura E. Jameson, Leslie A. Guerrero, Natalie E. Cain, Jessica Bolivar, Daniel A. Starr

**Affiliations:** Department of Molecular and Cellular Biology, University of California, Davis, Davis, CA, USA

## Abstract

KASH proteins in the outer nuclear membrane comprise the cytoplasmic half of LINC complexes that connect nuclei to the cytoskeleton. *Caenorhabditis elegans* ANC-1, an ortholog of Nesprin-1/2, contains actin-binding and KASH domains at opposite ends of a long spectrin-like region. Deletion of either the KASH or calponin homology (CH) domains does not completely disrupt nuclear positioning, suggesting neither KASH nor CH domains are essential. Deletions in the spectrin-like region of ANC-1 led to significant defects, but only recapitulated the null phenotype in combination with mutations in the trans-membrane span. In *anc-1* mutants, the ER was unanchored, moving throughout the cytoplasm, and often fragmented. The data presented here support a cytoplasmic integrity model where ANC-1 localizes to the ER membrane and extends into the cytoplasm to position nuclei, ER, mitochondria, and likely other organelles in place.

## Introduction

Cellular organization is an essential process. Organelles are interconnected and mostly constrained to specific subcellular locations when they are not actively transported longer distances by cytoskeletal motor proteins (van Bergeijk, Hoogenraad, & Kapitein, 2016). For example, nuclear positioning is essential for a wide variety of cellular and developmental processes, including fertilization, cell division, cell polarization, gametogenesis, central-nervous system development, and skeletal muscle function (Bone & Starr, 2016; Gundersen & Worman, 2013). Defects in nuclear positioning are associated with multiple neuromuscular diseases (Calvi & Burke, 2015; Folker & Baylies, 2013; Gundersen & Worman, 2013). Furthermore, the Golgi apparatus and centrosomes are often found next to the nucleus, while the ER and mitochondria are usually spread throughout the cell (van Bergeijk et al., 2016). The cellular tensegrity model posits that organelles are physically coupled to the cytoskeleton, plasma membrane, and extracellular matrix so that the cell acts as a single mechanical unit (Jaalouk & Lammerding, 2009; N. Wang, Tytell, & Ingber, 2009). The mechanisms that maintain cellular tensegrity and how it relates to organelle positioning are poorly understood, especially *in vivo*.

Nuclei are connected to the rest of the cell by LINC (linker of nucleoskeleton and cytoskeleton) complexes. SUN (Sad-1/UNC-84) proteins integral to the inner nuclear membrane and KASH (<u>K</u>larsicht, <u>A</u>NC-1, and <u>S</u>YNE <u>h</u>omology) proteins that span the outer nuclear membrane interact with each other in the perinuclear space to form LINC complexes. The cytoplasmic domains of KASH proteins interact with various components of the cytoskeleton (Luxton & Starr, 2014), while the nucleoplasmic domains of SUN proteins interact with lamins. Thus, LINC complexes bridge the nuclear envelope and mechanically couple the nucleoskeleton to the cytoskeleton (Chang, Worman, & Gundersen, 2015; Lee & Burke, 2018; Starr & Fridolfsson, 2010). LINC complex inhibition reduces the stiffness and increases the deformability of the entire cytoplasm in mammalian tissue culture cells, even far from the nuclear envelope and beyond the predicted reach of LINC complexes (Gill et al., 2019; Stewart-Hutchinson, Hale, Wirtz, & Hodzic, 2008). Whether LINC complexes maintain the mechanical properties of the cytoplasm *in vivo* is relatively unexplored.

Here, we investigate the role of LINC complexes with giant KASH proteins in organelle positioning in *Caenorhabditis elegans*. Most of the hypodermis of an adult *C. elegans* consists of a giant syncytium, the hyp7, containing 139 evenly-spaced nuclei that are anchored in place (Altun & Hall, 2002). Furthermore, *C. elegans* have invariant developmental lineages, are optically clear, and are easily genetically manipulated, making the hyp7 an ideal *in vivo* model to study organelle positioning. A LINC complex made of the KASH protein ANC-1 and the SUN protein UNC-84 is responsible for nuclear anchorage in *C. elegans* (Starr, 2019). UNC-84 is a canonical SUN protein that is orthologous to mammalian SUN1 and SUN2 (Malone, Fixsen, Horvitz, & Han, 1999) with a nucleoplasmic domain that interacts directly with the lamin protein LMN-1 (Bone, Tapley, Gorjanacz, & Starr, 2014). ANC-1 is an exceptionally large protein of up to 8545 residues with two tandem calponin-homology (CH) domains at its N terminus and a KASH domain at its C terminus (Starr & Han, 2002). ANC-1 orthologs *Drosophila* MSP-300 and mammalian Nesprin-1 Giant (G) and -2G have similar domain arrangements (Starr & Fridolfsson, 2010). Unlike MSP-300, Nesprin-1G, and -2G, which each contains greater than 50 spectrin-like repeats, ANC-1 consists of six tandem repeats (RPs) of 903 residues that are almost 100% conserved with each other at the nucleotide level (Liem, 2016; Rajgor & Shanahan, 2013; Starr & Han, 2002; Zhang et al., 2001). While spectrin-like repeats have not been identified in the ANC-1 RPs, most of ANC-1, is predicted to be highly helical, like spectrin (Starr & Han, 2002). The CH domains of ANC-1 interact with actin filaments *in vitro* and co-localize with actin structures *in vivo*, while the KASH domain of ANC-1 requires UNC-84 for its localization to the outer nuclear membrane (Starr & Han, 2002). UNC-84 is thought to interact with lamins to connect LINC to the nucleoskeleton while ANC-1 extends away from the outer nuclear membrane into the cytoplasm to tether nuclei to actin filaments (Starr & Han, 2002).

Evidence from multiple systems suggests that giant KASH orthologs might not solely function as nuclear tethers. We have observed that hyp7 syncytia in *anc-1* null animals display a stronger nuclear positioning defect than *unc-84* null animals (Cain et al., 2018; Jahed et al., 2019) and mitochondria are unanchored in *anc-1*, but not in *unc-84* mutants (Starr & Han, 2002). These results suggest that ANC-1 has LINC complex-independent roles for anchoring nuclei and mitochondria. Likewise, mitochondria and the ER, are mispositioned in *Drosophila msp-300* mutant muscles (Elhanany-Tamir et al., 2012). Finally, it remains to be determined if the CH domains of ANC-1 are necessary for nuclear anchorage. Mouse Nesprin-1 and -2 have isoforms lacking CH domains (Duong et al., 2014; Holt et al., 2016) and Nesprin-1 CH domains are dispensable for nuclear positioning during mouse skeletal muscle development (Stroud et al., 2017).

To address these ambiguities, we use ANC-1 to examine how giant KASH proteins position nuclei and other organelles. We find that for nuclear anchorage, the ANC-1 KASH domain plays a relatively minor role, and the CH domains are dispensable. Rather, multiple large cytoplasmic domains and the C-terminal transmembrane span of ANC-1 are required for nuclear anchorage. Moreover, in *anc-1* null mutants, the entire cytoplasm is disorganized, and the ER is unanchored and moves freely throughout the cytoplasm. Together, our results support a model in which ANC-1 associates with ER membranes to regulate the mechanical properties of the cytoplasm, thereby anchoring nuclei, mitochondria, ER, and other organelles.

## Results

### ANC-1 promotes proper nuclear anchorage through a LINC-complex-independent mechanism

Loss-of-function mutations in *anc-1* disrupt the even spacing of hyp7 syncytial nuclei (Cain et al., 2018; Starr & Han, 2002). We use the number of nuclei in contact with each other as a metric for hyp7 nuclear anchorage defects (Cain et al., 2018; Fridolfsson et al., 2018). In wild-type animals, very few hyp7 nuclei were touching. In contrast, over 50% of hyp7 nuclei were clustered with at least one other nucleus in *anc-1(e1873)* null mutants. Significantly fewer hyp7 nuclei were unanchored in *unc-84(n369)* null mutants (Figure 1C). This trend was also observed in adult syncytial seam cells (Figure S1).

**Figure 1.**
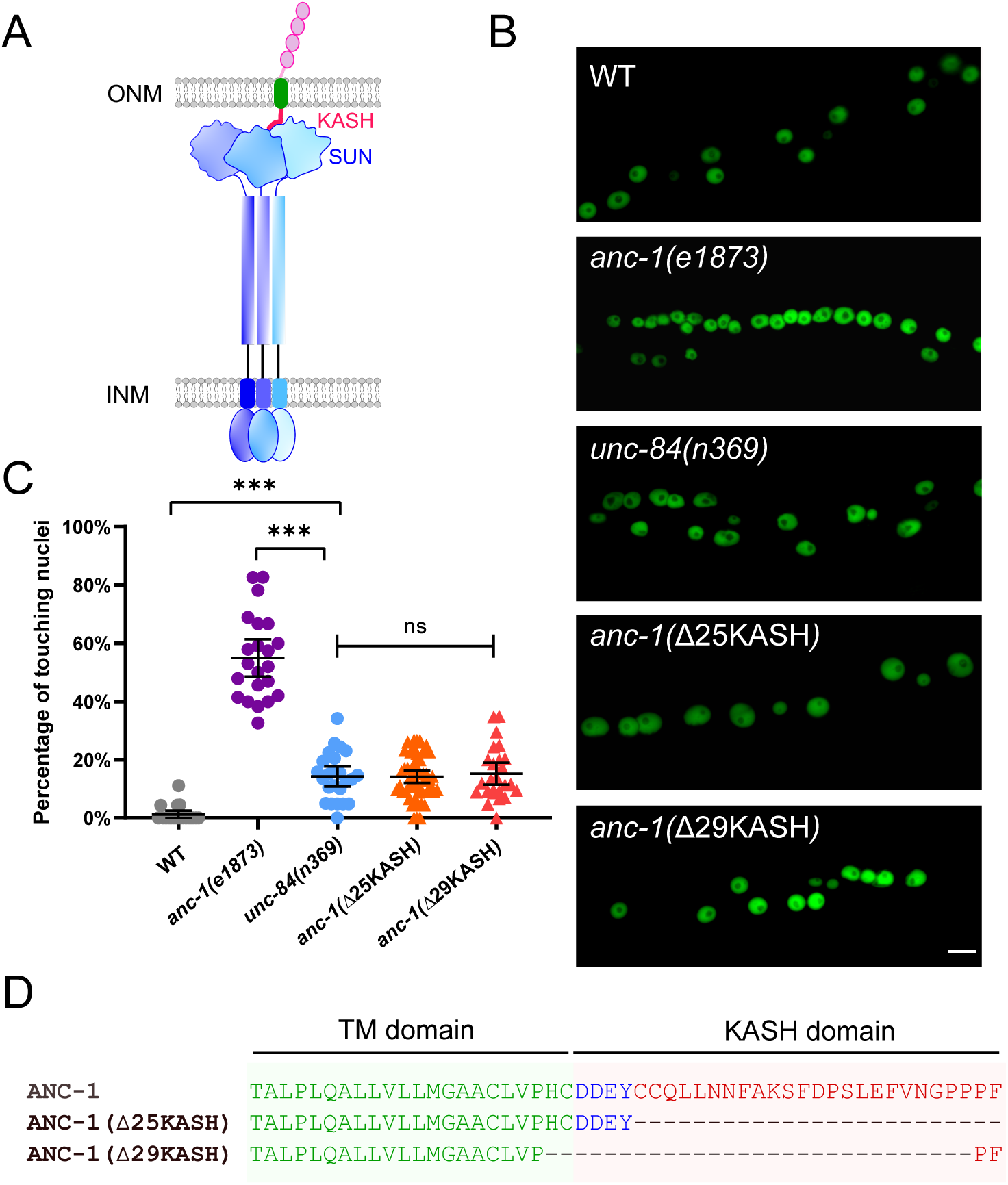
ANC-1 has a LINC complex -independent role in anchoring nuclei. (A) Model of the LINC complex. Trimers of the SUN protein UNC-84 (purple and blue) and the KASH protein ANC-1 (red, green and pink, only one of the trimers is shown) form the LINC complex, which spans the outer nuclear membrane (ONM) and inner nuclear membrane (INM). (B) Lateral views of adult *C. elegans* expressing hypodermal nuclear GFP in wild-type (WT) or indicated mutants. Scale bar, 10 µm. (C) Quantification of nuclear anchorage defects. Each point represents the percentage of touching nuclei on one side of a young adult animal. Means with 95% CI error bars are shown. Unpaired student two-tail *t*-test was used for statistical analysis. ns, not significant (p > 0.05); ***, *P* ≤ 0.001. n ≥ 20 for each strain. (D) Sequences of the transmembrane domain and the KASH domain of ANC-1 showing the deletions analyzed.

The greater severity of the nuclear anchorage defect in *anc-1* null mutants compared to *unc-84* suggests that ANC-1 plays additional roles in nuclear positioning independently of its SUN partner UNC-84. We used CRISPR/Cas9 gene-editing to delete the luminal peptides of the ANC-1 KASH domain (Figure 1D). We predicted the *anc-1(*ΔKASH) mutants would abrogate the interaction between ANC-1 and UNC-84 and phenocopy *unc-84*(*null*) animals. Two independent *anc-1(*ΔKASH) mutants exhibited mild nuclear anchorage defects similar to those observed in *unc-84(null)* mutants (Figure 1B-C). Together, these results suggest that the SUN/KASH interaction only partially contributes to nuclear anchorage, implicating the large cytoplasmic domain of ANC-1 as the major player in nuclear positioning.

### The ANC-1 N-terminal CH domains are not required for hyp7 nuclear positioning

We next deleted the CH domains at the N terminus of the largest isoforms of ANC-1, which are predicted to interact with actin, and replaced them with GFP using CRISPR/Cas9 gene editing. Hyp7 nuclei did not cluster in *anc-1(*ΔCH) mutants (Figure 2B). Thus, the CH domains of ANC-1 are not required for hyp7 nuclear anchorage.

**Figure 2.**
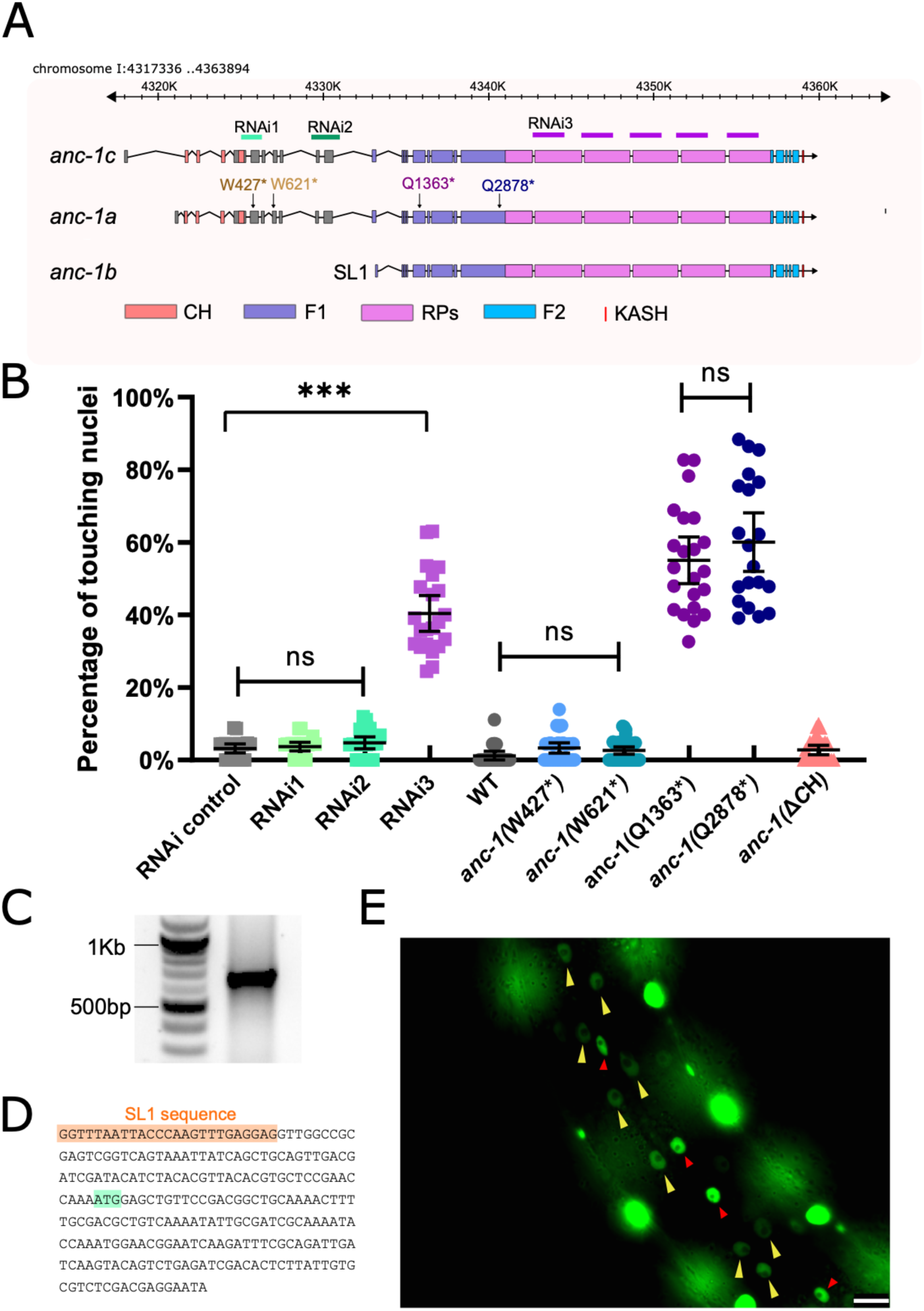
*anc-1b* is the major isoform in hyp7 nuclear anchorage. (A) Schematic gene structure of *anc-1a, b* and *c* isoforms (modified from the J-Brower in Wormbase). Domains are color-coded. The target regions of RNAi constructs are labeled. Premature stop mutations are indicated using the numbering the *anc-1a* isoform. (B) Quantification of nuclear anchorage defects in *anc-1* mutant and RNAi animals. Means with 95%CI are shown in the graph. Unpaired student two-tail *t*-test was used for statistical analysis. ns, not significant, p > 0.05; ***, *P* ≤ 0.001. n ≥ 20 for each strain. (C) An agarose gel showing the 5’-RACE products on the right lane. (D) Partial sequence of the 5’-RACE product. An SL1 sequence (orange) adjacent to the 5’ end of the *anc-1b* transcript was identified. The predicted start codon is in light green. (E) Lateral view of a worm showing the expression of nls::GFP driven by *anc-1b* promoter. Yellow arrows mark hyp7 nuclei, the red arrows mark seam cell nuclei and the bright, unmarked nuclei are in muscle cells. Scale bar is 10 µm.

Nesprin-1 and -2 have multiple splice isoforms, many of which are missing the CH domains (Rajgor, Mellad, Autore, Zhang, & Shanahan, 2012; Stroud et al., 2017). We hypothesized a shorter *anc-1* isoform lacking the CH domains would be sufficient for nuclear anchorage. RNAseq and expressed sequence tag data published on WormBase (Harris et al., 2020) suggest that *anc-1* has at least three isoforms (Figure 2A). We tested whether a shorter isoform lacking CH domains, *anc-1b*, is sufficient for hyp7 nuclear anchorage. RNAi constructs targeting the 5’ exons specific to the *anc-1a/c* long isoforms did not cause nuclear anchorage defects (Figure 2A-B). However, RNAi targeting a repetitive region in all three predicted isoforms, caused severe nuclear anchorage defects (Figure 2A-B). We analyzed four nonsense mutations and found that disrupting longer *anc-1a/c* isoforms did not result in significant nuclear anchorage defects. In contrast, both *anc-1(*Q1603*) and *anc-1(*Q2878*) alleles, which are predicted to add premature stop codons to all three predicted isoforms, led to severe, null-like nuclear anchorage defects (Figure 2A-B). These results suggest that the shorter *anc-1b* isoform lacking the CH domains is sufficient for nuclear anchorage.

We next tested whether *anc-1b* is expressed in hyp7. First, 5’ RACE (Rapid amplification of cDNA ends) was used to identify the start of the *anc-1b* predicted transcript (Figure 2C). The RACE product contained an SL1 sequence at its 5’ end, suggesting this represents the end of a bona fide transcript (Figure 2D). To test if the *anc-1b* isoform is expressed in hyp7, we fused the *anc-1b* promoter and ATG to an *nls::gfp::lacZ* reporter and expressed it in transgenic animals. The *anc-1b* promoter drove GFP expression in hyp7 (Figure 2E, yellow arrows). Taken together, these results suggest the conserved CH domains are not necessary for hyp7 nuclear anchorage and the *anc-1b* isoform expressed in the hypodermis plays a major role in hyp7 nuclear anchorage.

### Spectrin-like domains of ANC-1b are required for nuclear anchorage

We predicted that the six tandem repeats (RPs) of ANC-1 function analogously to the spectrin-like domains of Nesprin-1 and -2. We modeled the structure of pieces of the ANC-1 RPs using the protein-folding prediction software QUARK (Xu & Zhang, 2012, 2013) and found that they are predicted to form helical bundles remarkably similar to the structure of spectrin (Figure 3A, D) (Grum, Li, MacDonald, & Mondragon, 1999), suggesting the tandem repeats are analogous to spectrin-like domains.

**Figure 3.**
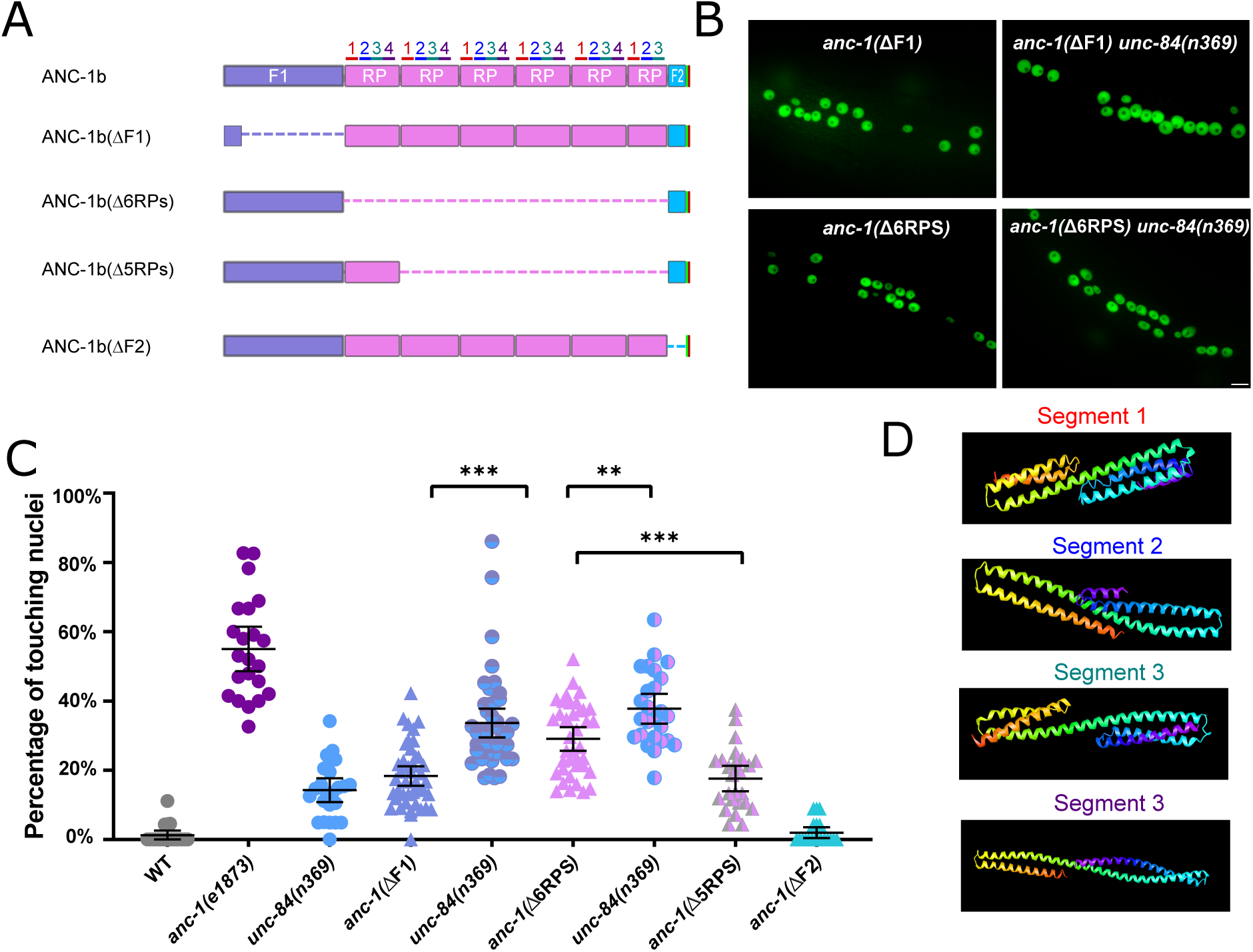
Cytoplasmic domain deletion analysis of ANC-1b. (A) Schematics of the ANC-1b cytoplasmic domain deletions. (B) Lateral views are shown of young adult *C. elegans* expressing hypodermal nuclear GFP in the indicated genotypes. Scale bar, 10 µm. (C) Quantification of nuclear anchorage in the wild-type as well as *anc-1b* domain deletion mutants. Each point represents the percentage of touching nuclei on one side of a young adult animal. Means with 95% CI error bars are shown. Unpaired student two-tail *t*-test was used for statistical analysis. **, *P* ≤ 0.01, ***, *P* ≤ 0.001. n ≥ 20 for each strain. (D) QUARK result for three fragments in the tandem repeats. The positions of the fragments are indicated in 3A.

We next tested the necessity of the ANC-1 spectrin-like repeats and the neighboring cytoplasmic domains for nuclear anchorage by making in-frame deletions of portions of *anc-1* using CRISPR/Cas9-mediated gene editing. The N-terminal fragment 1 (F1) contains 1969 residues of ANC-1b from the start codon to the start of the RPs and fragment 2 (F2) contains the 277 residues between the RPs and the C-terminal transmembrane span (Figure 3A). The deletion of the F1 domain or all six RPs caused severe nuclear anchorage defects (Figure 3 B-C). ANC-1b with only one of the normal six repeats had an intermediate nuclear anchorage defect that was significantly greater than the wild-type but less severe than the *anc-1(*Δ6RPS) mutant (Figure 3C). In contrast, the *anc-1(*ΔF2) mutant had no nuclear anchorage defect (Figure 3C). Since the *anc-1(*Δ6RPS) and *anc-1(*ΔF1) defects were less severe than null alleles, we made double mutants with *unc-84(n369)* to see if mutations in the cytoplasmic portions of ANC-1 were synergistic with mutations in the KASH domain. Both *anc-1(*ΔF1*); unc-84(n369)* and *anc-1(*Δ6RPS*); unc-84(n369)* double mutants significantly enhanced the nuclear anchorage defects of the single mutations (Figure 3C). However, the hyp7 nuclear anchorage defects in *anc-1(*ΔF1*); unc-84(n369)* and *anc-1(*Δ6RPS*); unc-84(n369)* double mutants were less severe than *anc-1(e1873)* null mutants, suggesting that multiple parts of ANC-1b mediate proper hyp7 nuclear positioning (Figure 3C). Together, these results indicate that 1) F1 and the RPs play roles in nuclear anchorage, 2) multiple repeats are necessary for normal function, and 3) the F2 region is dispensable for hyp7 nuclear positioning.

### The ER is unanchored in *anc-1* mutants

In addition to nuclear positioning, ANC-1 functions in mitochondria distribution and morphology in the hypodermis and muscle cells (Hedgecock & Thomson, 1982; Starr & Han, 2002). We therefore asked if ANC-1 also anchors other organelles. We characterized the ER in live hyp7 syncytia of *anc-1* and *unc-84* mutants using a single-copy GFP::KDEL marker. In wild type, the ER formed a branched network that was evenly distributed throughout hyp7 (Figure 4A). We used blind scoring to classify single images of each animal’s ER as normal (evenly distributed in what appeared to be sheets), mild defects (lots of sheet-like structures, but occasionally not uniformly spread throughout the syncytium), strong defects (considerable mispositioning and clustering of ER, but still in large units) or severe defects (complete mispositioning and extensive fragmenting of the ER) (Figure 4 and Supplemental Figure 2). About a third of wild-type adults had normally distributed ER, while the rest had mild ER positioning defects, perhaps due to the pressure on the worm from the coverslip (Figure 4A-B). However, ER networks in *anc-1* null mutants were severely disrupted and often fragmented (Figure 4A-B). We also observed the dynamics of the ER in live animals. In wild-type animals the hypodermal ER was anchored as an interconnected network and exhibited limited motion while the animal crawled (Figure 4C, E-F, Supplemental Movie 1). However, in *anc-1* null mutants, ER fragments drifted apart and often formed large aggregates, suggesting that the anchorage of the ER network was disrupted (Figure 4D-F, Supplemental Movie 2). To quantify this phenotype, we measured the change in distance between distinct points over time. In wild type, the average distance change between parts of the ER was less than 1 µm per second, whereas in *anc-1* mutants, the average change in distance more than doubled (Figure 4E), suggesting that the ER had lost its overall interconnectivity and that fragments were unanchored from the rest of the ER.

**Figure 4.**
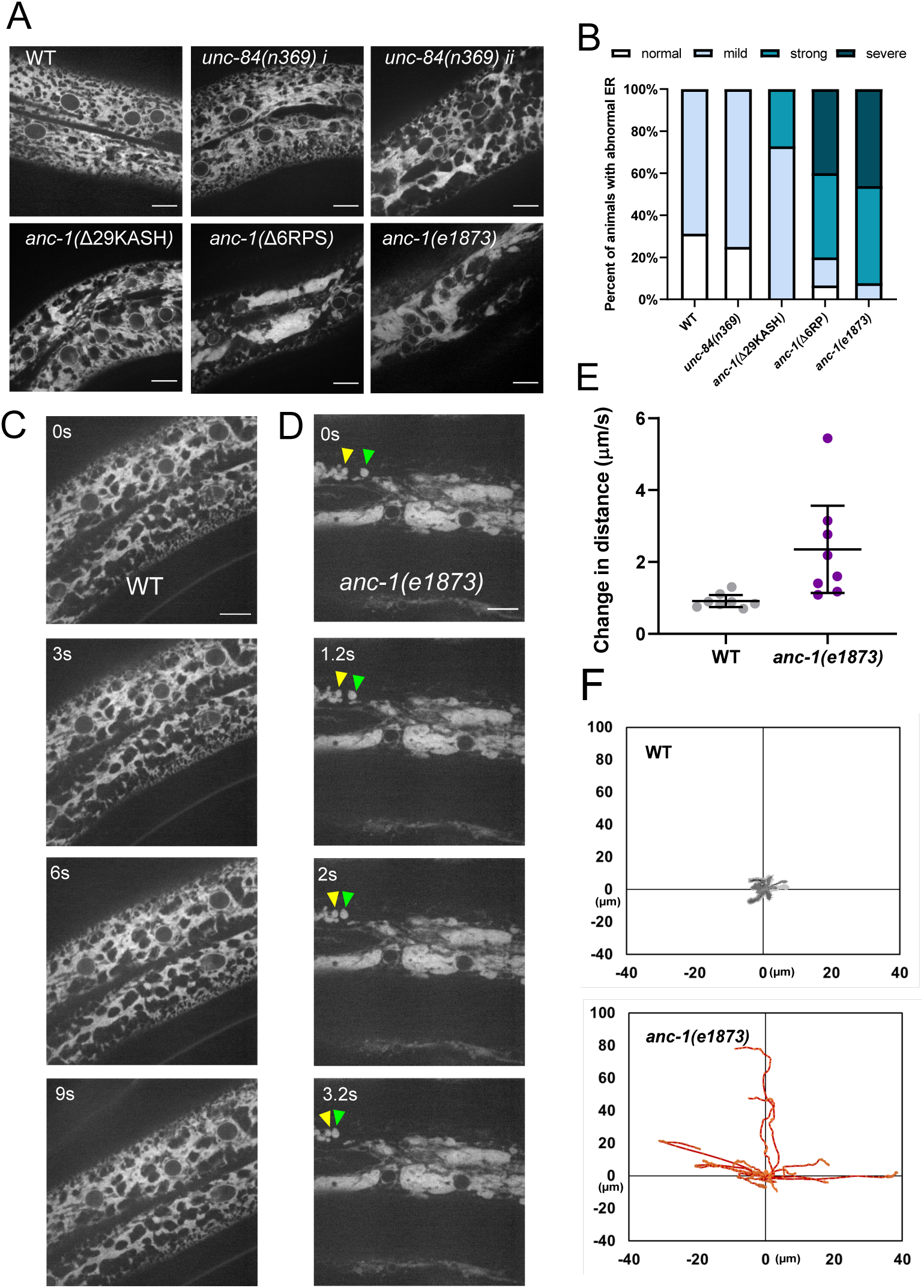
The ER is mispositioned in *anc-1* mutants. (A) Representative images of the hyp7 ER labeled with the GFP::KDEL marker in the young adult animals. (B) Scoring of the ER positioning defects. ER images of the listed strains were mixed and randomized for blind analysis by multiple researchers. n ≥ 11 for each strain. (C-D) Time-lapse images of hyp7 GFP::KDEL marker (C) over 9s in wild type or (D) over 3.2 seconds in *anc-1* null. Arrow heads show two fragments of ER that changed their relative distance from one another over a short period of time. (E-F) To quantify ER displacement, three spots on each WT and *anc-1(e1873)* movie were tracked. The average change in distance between two points in 200ms intervals is plotted in (E). The trajectories of the relative movements of each spot apart from the others are shown in (F). Eight movies of each strain were analyzed. Scale bar is 10 µm for all the images.

We next examined whether UNC-84 is required for ER positioning. Still images of *unc-84(n369)* null mutants scored blindly were similar to wild type ER (Figure 4A-B). However, in some videos of *unc-84(n369)* mutants, there were slight changes in the organization of the ER over time (Supplemental Movie 3), suggesting that *unc-84* null mutants had a minor ER positioning defect. Most *anc-1(*Δ29KASH) mutants had mild defects in ER positioning, and only about a quarter had more severe defects, significantly less than *anc-1(e1873)* null mutants. In contrast, more than 80% of *anc-1(*Δ6RPS) animals had strong or severe ER positioning defects, similar to *anc-1* null mutants (Figure 4A-B). These results indicated that ANC-1 is essential for ER positioning through mostly LINC complex-independent mechanisms.

### ANC-1 localizes to ER membranes

Since ANC-1 functions, in part, independently of LINC complexes at the nuclear envelope and because ANC-1 regulates ER positioning, we hypothesized that ANC-1 localizes to multiple membranes, including those away from the nuclear envelope. To study the localization of ANC-1, we tagged endogenous ANC-1 with GFP using CRISPR/Cas9 gene editing. The tag was placed either at the N-terminus of ANC-1b, or between the six tandem RPs and the F2 region to see if the opposite ends of ANC-1 localize to different structures (Figure 5A). Both strains were nearly wild type for hyp7 nuclear positioning (Figure 5B). Both GFP::ANC-1b and ANC-1::GFP::F2 localized in similar patterns throughout the cytoplasm of adult hyp7 syncytia (Figure 5C-D). To examine the localization of the ANC-1b isoform alone, we introduced a premature stop codon mutation to disrupt the longer *anc-1a* and *c* isoforms in the GFP::ANC-1b strain, which did not significantly change GFP::ANC-1b localization (Figure 5E). These data are consistent with our model that *anc-1b* plays the major role in hyp7 nuclear positioning.

**Figure 5.**
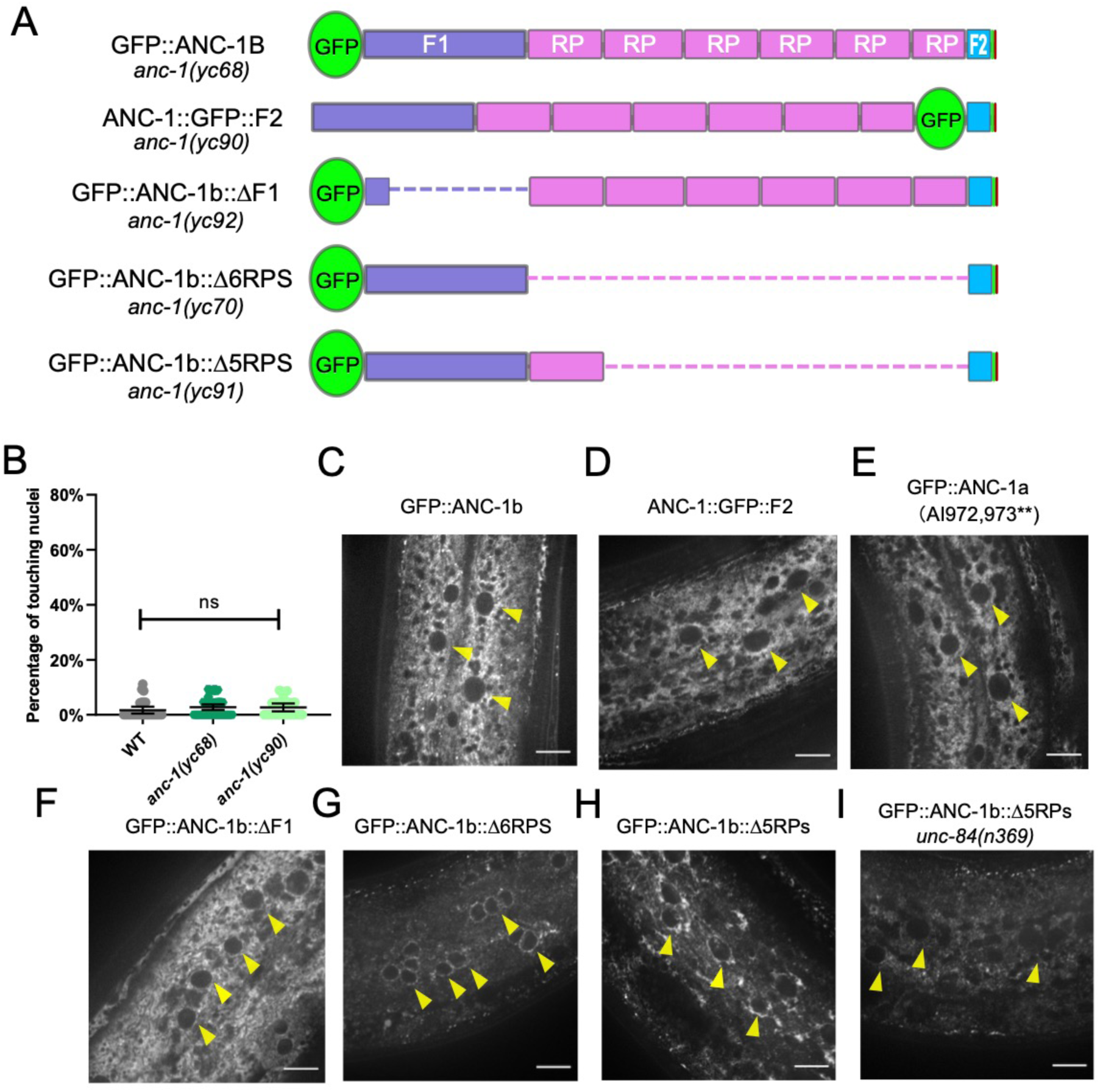
The subcellular localization of ANC-1. (A) Schematic depicting the ANC-1 GFP knock-in constructs with or without deletion of ANC-1 cytoplasmic domains. (B) Nuclear positioning in GFP::ANC-1b is wild type. Each point represents the percentage of touching nuclei on one side of a young adult animal. Means with 95% CI error bars are shown. n ≥ 20. (C-I) Confocal images of hyp7 subcellular localization in the indicated strains. Yellow arrow heads point to nuclei. Scale bar is 10 µm.

We next examined whether the cytoplasmic domains of ANC-1b are required for localization. Deletion of the F1 domain did not dramatically change localization of GFP::ANC-1b (Figure 5F). However, deletion of the six tandem repeats enriched GFP::ANC-1b around the nuclear envelope (Figure 5G), as did the deletion of five of the six repeats (Figure 5H). The nuclear envelope enrichment of GFP::ANC-1b(Δ5RPs) was not observed when *unc-84* was mutated (Figure 5I), suggesting the nuclear envelope enrichment of GFP::ANC-1b is UNC-84-dependent.

The ANC-1 localization pattern in Figure 6 is consistent with a model where ANC-1 localizes to ER membranes. In this model, a construct containing the transmembrane span near the C terminus of ANC-1 but lacking the C-terminal KASH domain would be sufficient for ER localization. Deletion of both the transmembrane span and the luminal KASH domain with CRISPR/Cas9 resulted in a significantly more severe nuclear anchorage defect than deleting only the luminal KASH domain (Figure 6B). Similarly, ER positioning defects were worse in *anc-1(*ΔTK) than in *anc-1(*Δ25KASH) animals (Figure 6C). We next examined the role of the neck region, which is located immediately adjacent to the cytoplasmic side of the transmembrane span. When GFP was knocked-in between the neck region and the transmembrane span to make *anc-1(yc36[anc-1::gfp3Xflag::kash])*, it caused a significant nuclear anchorage defect (Figure 6A-B). Moreover, extending the deletion in *anc-1(*ΔF2), which had no hyp7 nuclear anchorage defect (Figure 3C), an additional 9 residues to remove the neck, caused a significant nuclear anchorage defect (Figure 6A-B). This suggests that the neck region next to the transmembrane span of ANC-1 plays a role in nuclear positioning, perhaps by targeting the C terminus of ANC-1 to a membrane.

**Figure 6.**
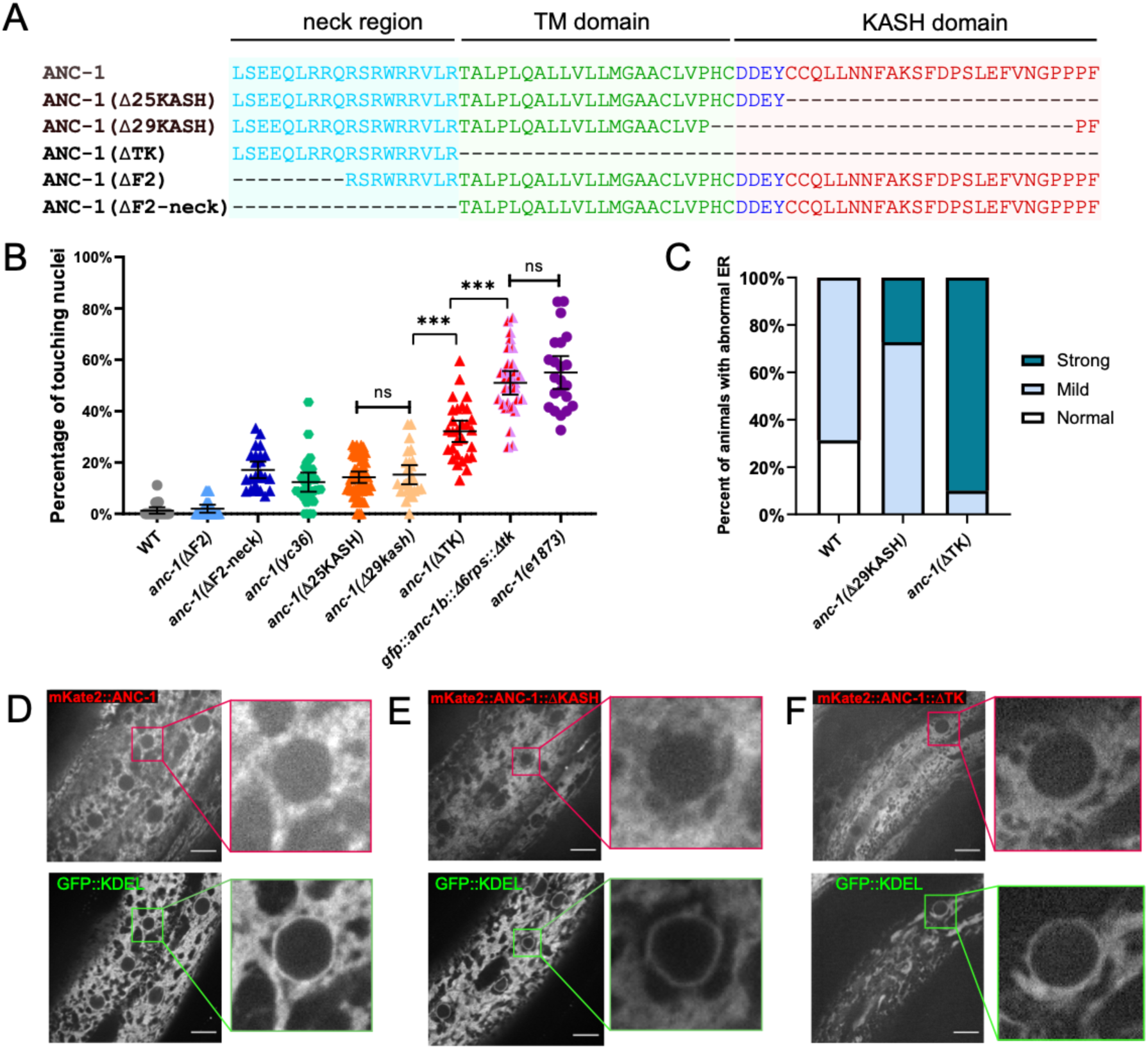
The transmembrane domain of ANC-1 targets it to the ER. (A) Sequence of deletion in the neck region (blue), transmembrane (TM) span (green) or the KASH domain (red). (B) Quantification of nuclear anchorage defects in *anc-1* mutants. Each point represents the percentage of touching nuclei on one side of a young adult animal. Means with 95% CI error bars are shown. n ≥ 20. (C) Qualitative analysis of the ER anchorage defects as in Figure 4. Significantly more *anc-1(*ΔTK) animals show strong ER anchorage defects than *anc-1(*Δ29KASH) mutants (p ≤ 0.01 by Fisher’s exact test). Sample sizes were all ≥ 10. (D-F) The co-localization of the GFP::KDEL ER marker with mKate2::ANC-1 (D), mKate2::ANC-1:: Δ25KASH (E) and mKate2::ANC-1:: ΔTK (F). Zoomed-in views of the boxed regions are shown on the right of each lane. Scale Bar is 10 µm.

As shown above, a double mutant that lacks both the ANC-1 repeat region and LINC complex function in the form of an *unc-84(null)* was not as severe as an *anc-1(null)* mutant (Figure 3B). This suggests that a third domain of ANC-1, in addition to the KASH and RPs, is involved in positioning nuclei. To test this hypothesis, we made an *anc-1(gfp::anc-1b::Δ6rps::Δtk)* mutant line and found that it had a severe nuclear anchorage defect that was not significantly different from the *anc-1(e1873)* null mutant (Figure 6B). Together, these data support a model in which the transmembrane span is important for targeting ANC-1 to the ER/nuclear envelope membrane where ANC-1 then positions the ER and nuclei via LINC complex-independent mechanisms.

To further examine the role of the transmembrane span in ER and nuclear positioning, we observed the co-localization of ANC-1 mutant fluorescent fusion proteins with an ER marker. An mKate2::ANC-1b fusion protein had a similar distribution to GFP::KDEL (Figure 6D), although it was less intense at the nuclear envelope. The slight discrepancy between localization of mKate2::ANC-1b and GFP::KDEL could be explained by the distance between the ANC-1 N terminus and the ER membrane. Deleting the KASH domain from mKate2::ANC-1b (mKate2::ANC-1b::ΔKASH) did not significantly change its localization pattern relative to the wild-type construct (Figure 6E). In contrast, deletion of both the KASH peptide and the transmembrane span resulted in more diffuse localization, often in areas of the cytoplasm that lacked ER (Figure 6F). Together, these results suggest that 1) ANC-1b normally localizes to the cytoplasmic surface of the ER and 2) that the transmembrane span, but not the KASH domain, is required for ER localization. This localization pattern is consistent with the above findings that *anc-1(*ΔTK) mutants had more severe nuclear and ER positioning defects than *anc-1(*Δ25KASH) mutants.

### Depletion of ANC-1 disrupts nuclear morphology and causes developmental defects

Nuclear shape changes were observed during live imaging in *anc-1* mutants consistent with a model where *anc-1* mutant nuclei are susceptible to pressures from the cytoplasm, perhaps crashing into lipid droplets that corresponded with dents in nuclei. We therefore quantified nuclear size and shape in *anc-1* mutants to better characterize how nuclear and/or ER movements affect the nuclear structure. Adult syncytial hyp7 nuclei were significantly smaller in *anc-1(*Δ6RPS) and *anc-1(e1873)* mutants compared to wild-type (Figure 7A). Furthermore, the shape of *anc-1(e1873)* hyp7 nuclei, as measured by circularity and solidity, were significantly less round than wild type (Figure 7B).

**Figure 7.**
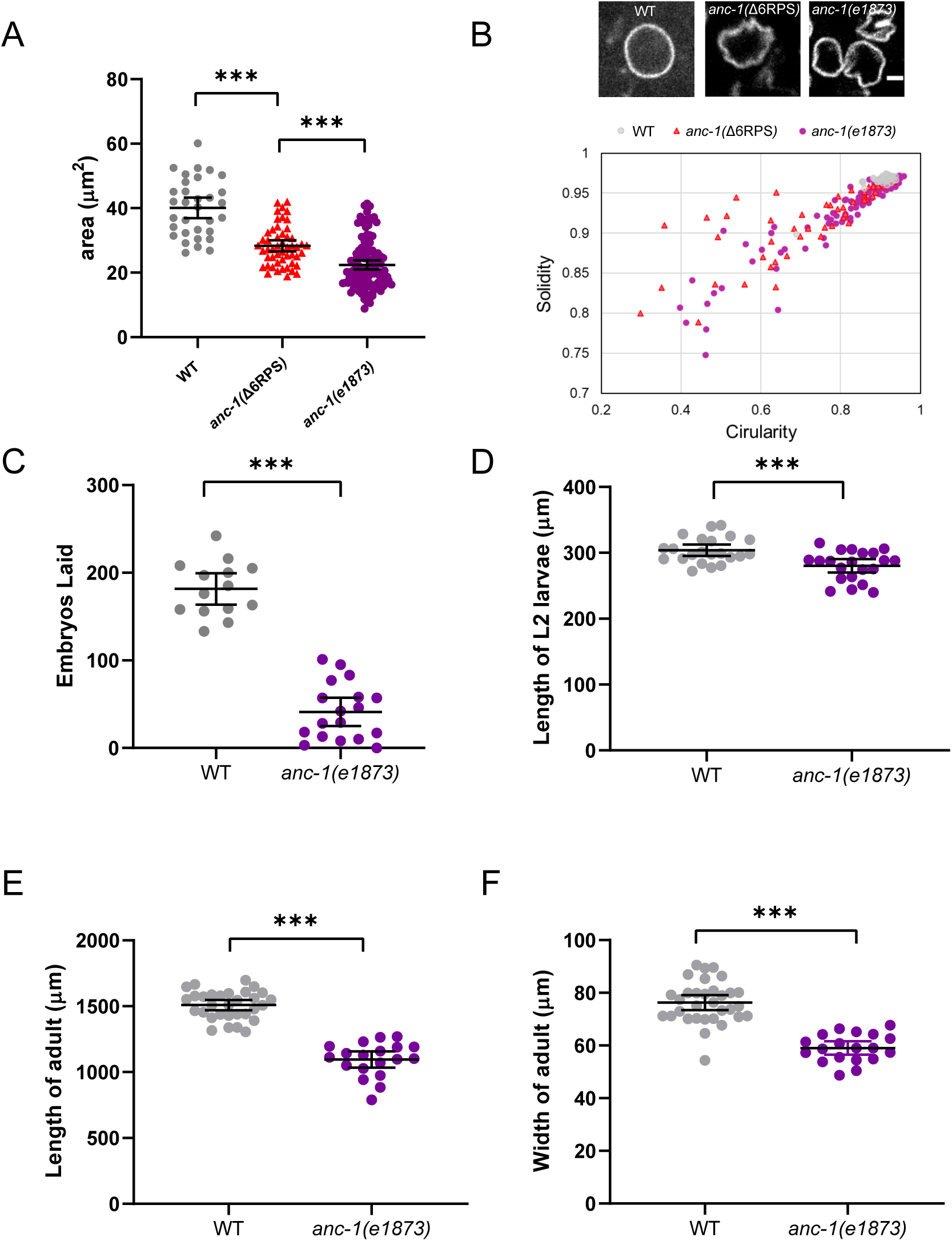
*anc-1* mutants have developmental defects. (A) The area of cross-sections of hyp7 nuclei is shown. Each dot represents the area of a single nucleus. n = 32 for WT, n = 50 for *anc-1(*Δ6RPS), n = 111 for *anc-1(e1873)*. (B) Top panel: Representative images of hyp7 nuclei marked by EMR-1::mCherry of WT, *anc-1(*Δ6RPS) and *anc-1(e1873)* mutants. Scale bar, 2µm. Bottom panel: Plot of the solidity and the circularity. (C-F) The brood size (C), length of the L2 larvae (D), adult length (E) and width (F) are significantly reduced in *anc-1(e1873)* mutants. Each dot represents a single animal. n ≥ 14 for (C) and n ≥ 19 for (D-F). Means with 95% CI error bars are shown. Unpaired student two-tail t-test was used for statistical analysis. **, *P* ≤ 0.01, ***, *P* ≤ 0.001.

Given the nuclear anchorage, ER positioning, and nuclear shape defects observed in *anc-1* mutants, we examined whether these animals might have other developmental or growth defects. Despite producing fertile adults, *anc-1* null animals had severe developmental defects. The brood size of *anc-1(e1873)* mutants was less than 25% of wild type (Figure 7C) and *anc-1(e1873)* mutants had significantly smaller body sizes throughout larval and adult stages (Figure 7D-F). Together these results suggest that there are developmental consequences associated with organelle positioning defects in *anc-1* mutants.

## Discussion

LINC complexes consisting of giant KASH proteins (*C. elegans* ANC-1, *Drosophila* MSP300, and mammalian Nesprin-1 and -2) and canonical SUN proteins (*C. elegans* UNC-84, *Drosophila* Koi, and mammalian Sun1 and Sun2) are thought to anchor nuclei by tethering them to the cytoskeleton (Starr & Fridolfsson, 2010). ANC-1 was thought to connect actin filaments to the nuclear envelope using conserved CH and KASH domains at its N and C termini, respectively (Starr & Han, 2002). However, results here and elsewhere indicate that this model is not sufficient to explain the role of KASH proteins in the cell. We propose a cytoplasmic integrity model, where giant KASH proteins function, largely independently of LINC complexes at the nuclear envelope and through mechanisms that do not require their CH domains. We find ANC-1 also localizes close to the ER throughout the cell. The large cytoplasmic domains of ANC-1 are required for positioning nuclei, ER, and likely other organelles. In the absence of ANC-1, the contents of the cytoplasm are disconnected from each other, unanchored in place, and sometimes fragmented. The cytoplasm and organelles freely flow throughout the hypodermal syncytia as worms crawl. It appears that ANC-1 is required to organize the entire cytoplasm.

Our finding that ANC-1 works independently of LINC complexes during hyp7 nuclear positioning has significant implications for how the field currently understands the role of LINC complexes in development and disease. A dominant negative approach relies on overexpression of KASH domains to displace endogenous KASH proteins from the nuclear envelope (Grady, Starr, Ackerman, Sanes, & Han, 2005; Starr & Han, 2002; Tsujikawa, Omori, Biyanwila, & Malicki, 2007). Alternatively, a mini-nesprin-2 construct, consisting of the calponin homology and KASH domains with a few spectrin repeats, is commonly used in rescue experiments (Davidson et al., 2020; Luxton, Gomes, Folker, Vintinner, & Gundersen, 2010; Ostlund et al., 2009). If ANC-1 and other giant KASH orthologs have major LINC complex independent functions, these approaches might have more caveats than previously thought.

We found that the CH domain of ANC-1 is dispensable for ER and nuclear positioning. Similar findings suggest the Nesprin-1 or -2 CH domains are also dispensable for the development of mouse skeletal muscles and the epidermis (Luke et al., 2008; Stroud et al., 2017). Moreover, the CH domains are not required for Nesprin-2 mediated neuronal migration in the developing rat brain (Goncalves, Quintremil, Yi, & Vallee, 2020). Thus, the significance of the conserved CH domain is not clear. The CH domain of Nesprin-2 is required to form transmembrane actin-associated lines on the nuclear envelope during rearward nuclear movement in mouse NIH3T3 fibroblasts polarizing for migration and to accumulate Nesprin-2 at the front of nuclei in cultured mouse embryonic fibroblasts migrating through constrictions (Davidson et al., 2020; Luxton et al., 2010).

Most of the other ANC-1 domains, including the C-terminal transmembrane span and the large cytoplasmic repeat regions, are required for nuclear anchorage. Portions of the tandem repeats may be arranged in bundles of three helices, reminiscent of spectrin repeats (Liem, 2016). Thus, we hypothesize that ANC-1, like the other giant KASH orthologs MSP300 and Nesprin-1 and -2, consists largely of spectrin repeats with a C-terminal transmembrane domain that attaches ANC-1 to the contiguous ER and outer nuclear membrane.

In general, giant KASH proteins localize strongly to the nuclear envelope in multiple systems (Starr & Fridolfsson, 2010). Evidence for giant KASH protein localization to other subcellular locations have been largely ignored or attributed to over-expression artifacts (Zhang et al., 2001) or the loss of KASH protein function (Roux et al., 2009). Antibodies against ANC-1 mostly localize away from nuclei and are only clearly enriched at the nuclear periphery in the somatic gonad (Starr & Han, 2002) and Nesprin-1 localizes to ciliary rootlets (Potter et al., 2017). Yet, most models, including the nuclear tethering model, focus primarily on nuclear envelope localization. We found that GFP::ANC-1b was not enriched at the nuclear envelope in most tissues. Instead, our findings suggest that ANC-1 functions throughout the cytoplasm with its C-terminus in the ER membrane.

We observed that the ER is severely mispositioned in *anc-1* null mutant hypodermal cells. Also, mitochondria are misshaped and mispositioned in the *anc-1* null mutants, but not in *unc-84* nulls (Hedgecock & Thomson, 1982; Starr & Han, 2002). Two different mechanisms could explain how nuclei, mitochondria, and the ER are all mispositioned in *anc-1* mutants. First, abnormal organelle clusters could result from a failure in the active positioning process (Folker & Baylies, 2013; Roman & Gomes, 2018). Alternatively, organelles could lose physical connections to the rest of the cytoplasm, resulting in organelles being passively pushed around in response to external mechanical forces, such as those generated when the worm crawls. Live imaging favors the second model. As *anc-1* mutant worms bent their bodies, ER fragments rapidly moved in a bulk flow of cytoplasm along with nuclei. The nuclei became misshapen and the ER often fragmented. Nuclei and the ER appeared to be completely detached from any sort of cytoplasmic structural network in *anc-1* mutants. There is precedent that giant KASH proteins could mechanically stabilize the cytoplasm. In cultured mouse NIH3T3 fibroblasts, disrupting KASH proteins leads to the decreased stiffness across the entire cell, as determined by single-particle microrheology experiments (Stewart-Hutchinson et al., 2008). Therefore, we propose that ANC-1 and other giant KASH proteins normally function at the ER periphery as important crosslinkers to maintain the mechanical integrity of the cell.

ANC-1 likely maintains the mechanical integrity of the cytoplasm via cytoskeletal networks. Disruption of MSP300 leads to disruption of both actin and microtubule networks in *Drosophila* muscles (S. Wang, Reuveny, & Volk, 2015). Giant KASH proteins can interact with actin independently of their CH domains. ANC-1 binds to AMPH-1/BIN1, which is involved in actin nucleation (D’Alessandro et al., 2015), and Nesprin-2 binds to the actin regulators Fascin and FHOD1 (Jayo et al., 2016; Kutscheidt et al., 2014). Nesprin-1 interacts with Akap450 to form microtubule organizing centers at the nuclear envelope of mouse muscles (Elhanany-Tamir et al., 2012; Gimpel et al., 2017; Zheng et al., 2020). Intermediate filaments and spectraplakins have also been implicated in nuclear positioning (Ralston et al., 2006; S. Wang et al., 2015; Zheng et al., 2020). How ANC-1b might directly interact with cytoskeletal components requires further investigations.

In summary, we propose a cytoplasmic integrity model, for how ANC-1 and giant KASH orthologs position nuclei and other organelles. First, ANC-1 is targeted to the ER through its C-terminal transmembrane span. The large cytoplasmic domain of ANC-1 likely consists of divergent spectrin-like repeats that could serve as elastic filaments. ANC-1 filaments are predicted to interact with various components of the cytoskeleton, allowing ANC-1 to serve as a cytoskeletal crosslinker to maintain the mechanical integrity of the cytoplasm. Eliminating ANC-1 abolishes interconnectivity between cytoskeletal networks and organelles become mis-positioned. Extracellular mechanical pressure that is generated as the worm bends, leads to rapid cytoplasmic fluid flows and positioning defects in nuclei, ER, mitochondria, and likely other organelles. In this model, ANC-1 maintains cellular connectivity across the cytoplasm by anchoring organelles in place.

## Supporting information

Sopplemental Movie 3

## Methods

### *C. elegans* genetics

*C. elegans* strains were maintained on nematode growth medium plates seeded with OP50 *E. coli* at the room temperature (approximately 22°C) (Brenner, 1974). All the strains used in this study are listed in Table 1. Some strains, including N2, which was used as wild type, were obtained from the Caenorhabditis Genetics Center, funded by the National Institutes of Health Office of Research Infrastructure Programs (P40 OD010440). Strains VC40007, VC20178 and VC40614 were provided by the *C. elegans* Reverse Genetics Core Facility at the University of British Columbia (Thompson et al., 2013). Strain RT3739 (*pwsi83*) was generously provided by Barth Grant (Rutgers University, NJ, USA). UD522 (*ycEx249[pcol-19::gfp::lacZ, pmyo-2::mCherry*]) was previously described (Cain et al., 2018). Male strains of RT3739, UD522 and BN147 were made to cross into *anc-1* or *unc-84* mutants. Alternatively, *pcol-19::gfp::lacZ* was introduced into some mutants by standard germ-line transformation to make UD736 and UD737 (Cain et al., 2018; Mello, Kramer, Stinchcomb, & Ambros, 1991).

**Table 1:**
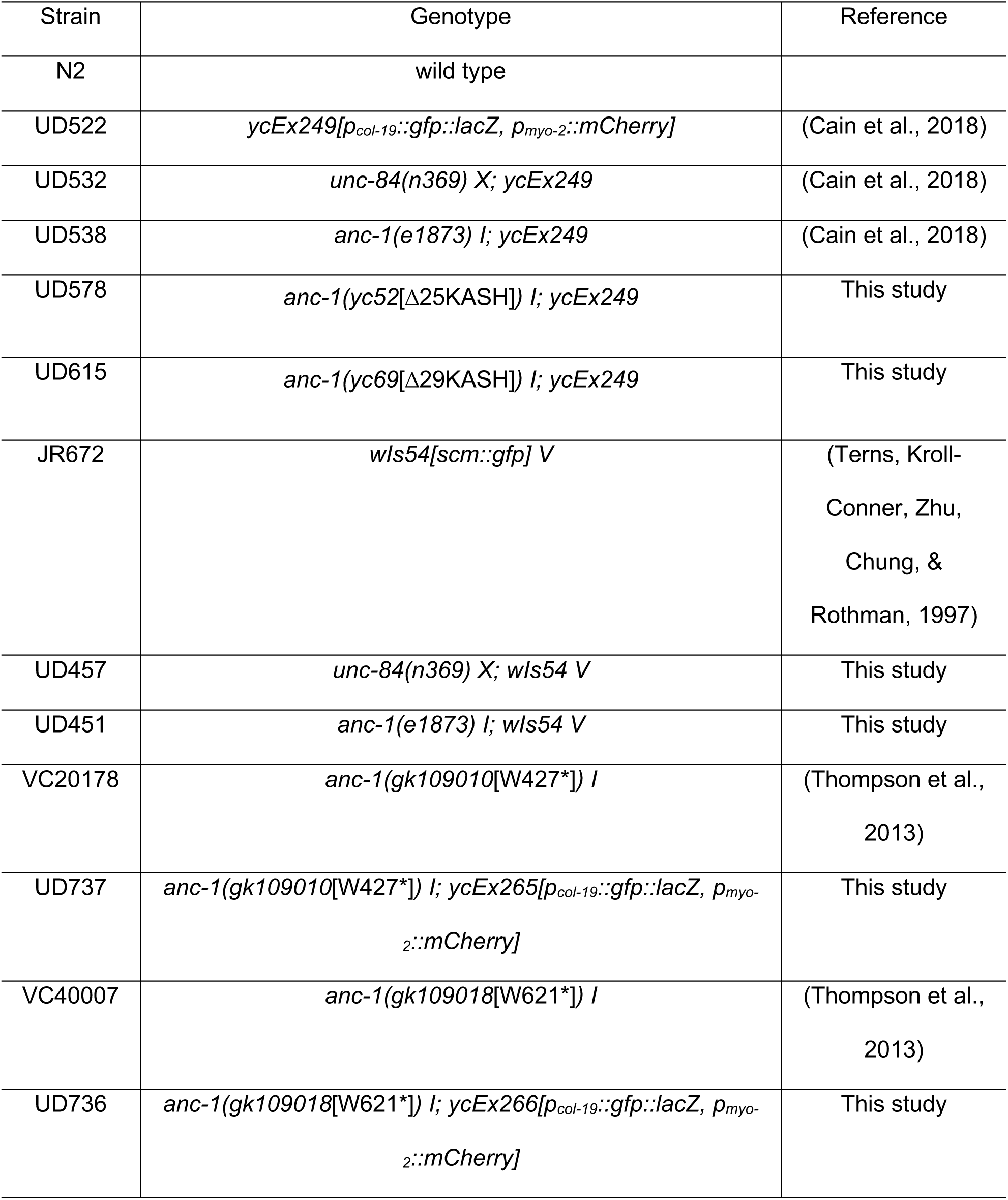
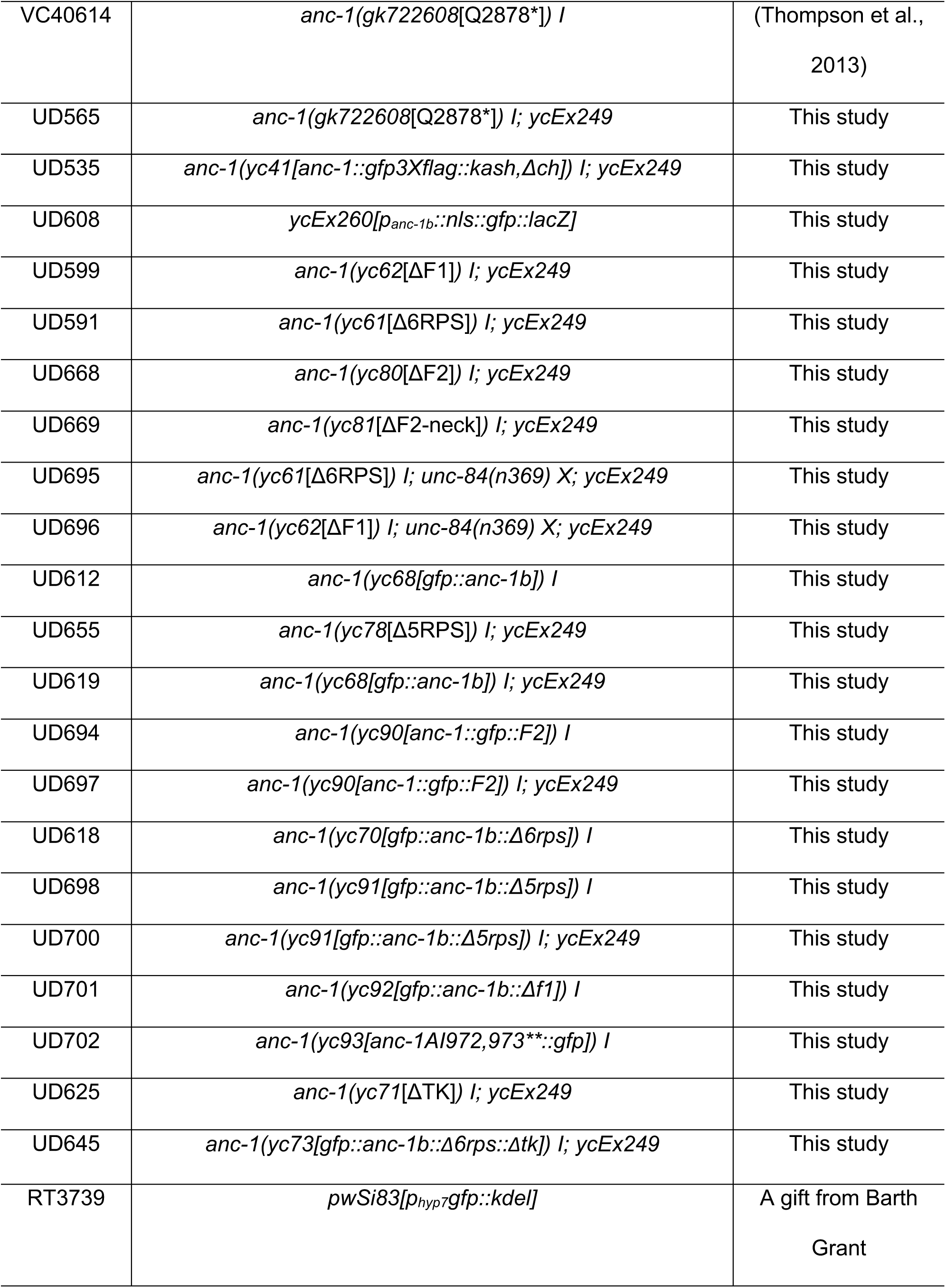
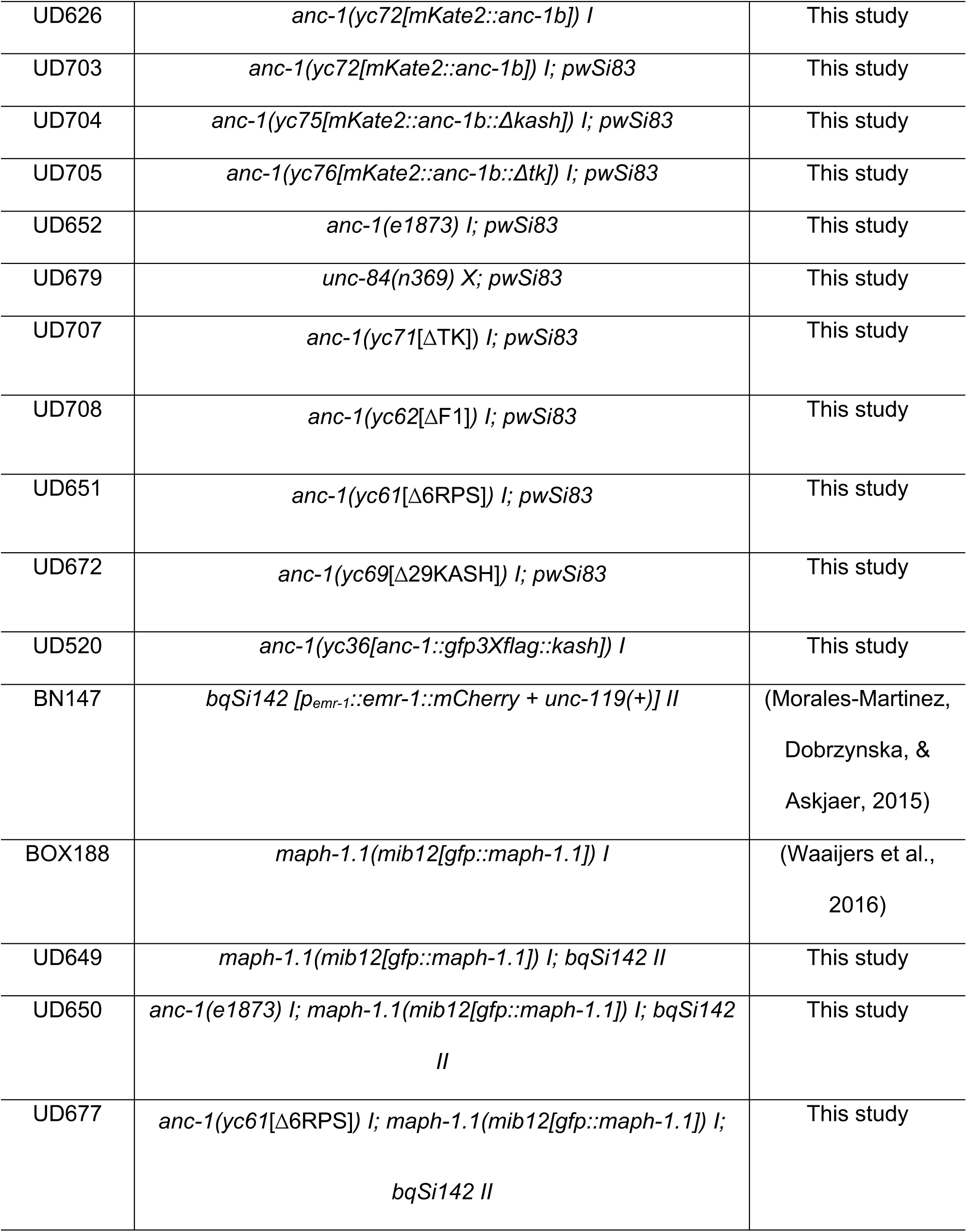
*C. elegans* Strains in this study.

For *anc-1* RNAi feeding experiments, L4 stage animals were transferred onto NGM plates seeded with bacteria expressing dsRNA (Timmons & Fire, 1998). The clones from the Ahringer RNAi library (Source Bioscience) (Kamath & Ahringer, 2003) were confirmed by Sanger sequencing.

Nuclear anchorage assays were performed as described (Cain et al., 2018; Fridolfsson et al., 2018). Briefly, L4 worms with GFP-marked hypodermal nuclei were picked onto fresh plates 20 hours before scoring. Young adults were mounted on 2% agarose pads in ∼5 µl of 1mM tetramisole in M9 buffer (Fridolfsson et al., 2018). Syncytial hyp7 nuclei were scored as touching if a nucleus was in contact with one or more neighboring nuclei. Only one lateral side of each animal was scored.

For the brood size assay, starting at the L4 stage, single animals were transferred onto fresh OP50 *E. coli* plates every 24 hours for seven days. The number of embryos laid were counted immediately after the removal of the animal each day.

To measure body size, embryos laid within an hour were collected and cultured for 24 hours and 69 hours to reach the L2 stage and the adult stage, respectively. Animals were mounted on 2% agarose pads in ∼5 µl of 1 mM tetramisole in M9 buffer for imaging.

### 5’-Rapid amplification of cDNA ends (5’-RACE)

Total RNA was extracted from mixed stages of *C. elegans* using the RNeasy kit (QIAGEN). First-strand cDNAs were generated with the ThermoScript RT-PCR system using an *anc-1* antisense oligonucleotide (ods2572: 5’-ATAGATCATTACAAGATG-3’). Purification and TdT tailing of the first-strand cDNA were performed by the 5’ RACE System for Rapid Amplification of cDNA Ends, version 2.0 (Invitrogen, Cat. No. 18374058). The target cDNA was amplified PCR using the provided 5’ RACE Abridged Anchor Primer and an *anc-1* specific primer: ods2574 (5’-GTCGGCGTCTGAAGGAAAGA-3’). The PCR product was purified using the QIAquick PCR purification kit (QIAGEN) and Sanger sequencing was performed by Genewiz.

### Plasmid construction and transformation with extrachromosomal arrays

To generate plasmid *p*_*anc-1b*_::*nls::gfp::lacZ* (pSL835), a 2.56 kb fragment of genomic DNA upstream of the start codon of *anc-1b* was amplified with primers ods2491 (5’-TACCGAGCTCAGAAAAAATGACTGTGAGTATAGTCATTTTCCGCT-3’) and ods2492 (5’-GTACCTTACGCTTCTTCTTTGGAGCCATTTTGGTTCGGAGCAC-3’) to replace the *col-19* promoter of *p*_*col-19*_::*gfp::lacZ* (pSL779) (Cain et al., 2018). N2 animals were injected with 45 ng/µl of pSL835, 50ng/µl of pBluescript SK, and 2.5 ng/µl of pCFJ90 (*p*_*myo-2*_::*mCherry*) (Frokjaer-Jensen et al., 2008) by standard *C. elegans* germline transformation (Evans, 2006) to make strain UD608 (*ycEx260[Panc-1b::nls::gfp::lacZ-2]*).

### CRISPR/Cas9 mediated gene editing

Knock-in strains were generated using a *dpy-10* Co-CRISPR strategy (Arribere et al., 2014; Paix, Folkmann, Rasoloson, & Seydoux, 2015; Paix, Folkmann, & Seydoux, 2017). All crRNA and repair template sequences are in Table 2. An injection mix containing 0.12 µl *dpy-10* crRNA (0.6 mM) (Horizon Discovery/Dharmacon), 0.3 µl target gene crRNA (0.6mM) for one locus editing or 0.21 µl of each crRNA (0.6mM) for multi-loci editing and 1.46 µl (one locus) or 1.88 µl (two loci) universal tracrRNA (0.17mM) (Horizon Discovery/Dharmacon) precomplexed with purified 7.6 µl of 40 µM Cas9 protein (UC Berkeley QB3) and 0.29 µl of the *dpy-10* single-strand DNA oligonucleotide (ssODN) (500ng/µl) repair templates and 0.21 µl ssODN repair template (25 µM) for the target gene editing or up to 500 ng double-strand DNA were injected to the germline of the hermaphrodite young adults. For *anc-1(yc52*[Δ25KASH]) *I, anc-1(yc69*[Δ29KASH]*) I, anc-1(yc62*[ΔF1]*) I, anc-1(yc61*[Δ6RPS]*) I, anc-1(yc80*[ΔF2]*) I, anc-1(yc78*[Δ5RPS]*) I, anc-1(yc91[gfp::anc-1b::Δ5rps]) I, anc-1(yc70[gfp::anc-1b::Δ6rps]) I, anc-1(yc92[gfp::anc-1b::Δf1]) I, anc-1(yc93[anc-1AI972,973**::gfp]) I*, single-strand DNA (SSD) (synthesized by Integrated DNA Technologies, IDT) was used as repair template. For GFP and mKate2 knock-in strains, double-strand DNA repair templates were amplified with PCR from the plasmids pSL779 for *gfp* or pSL780 for mKate2 using Phusion polymerase and the primers listed in table 2 (New England Biolabs) (Bone, Chang, Cain, Murphy, & Starr, 2016; Cain et al., 2018).

**Table 2:**
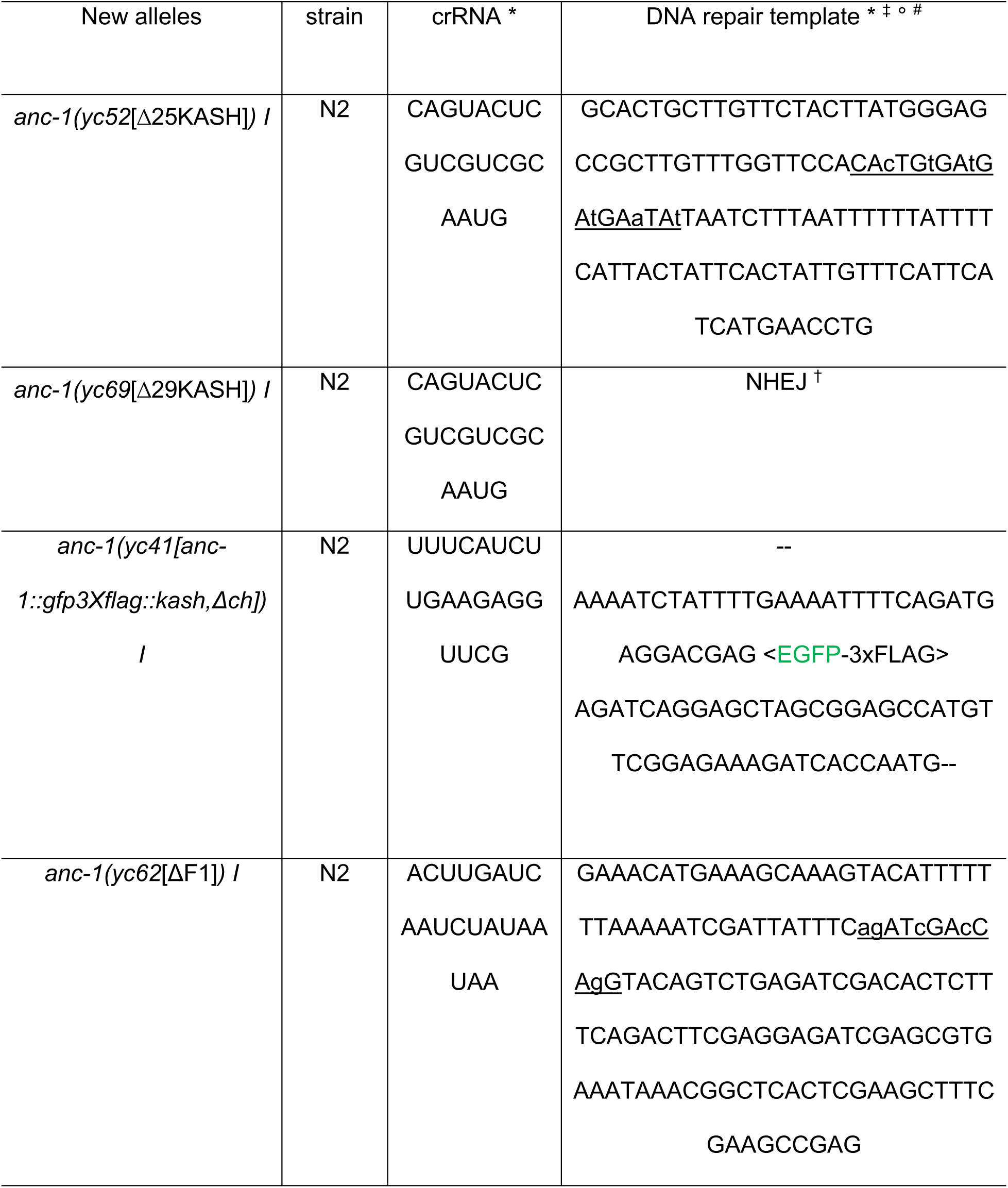
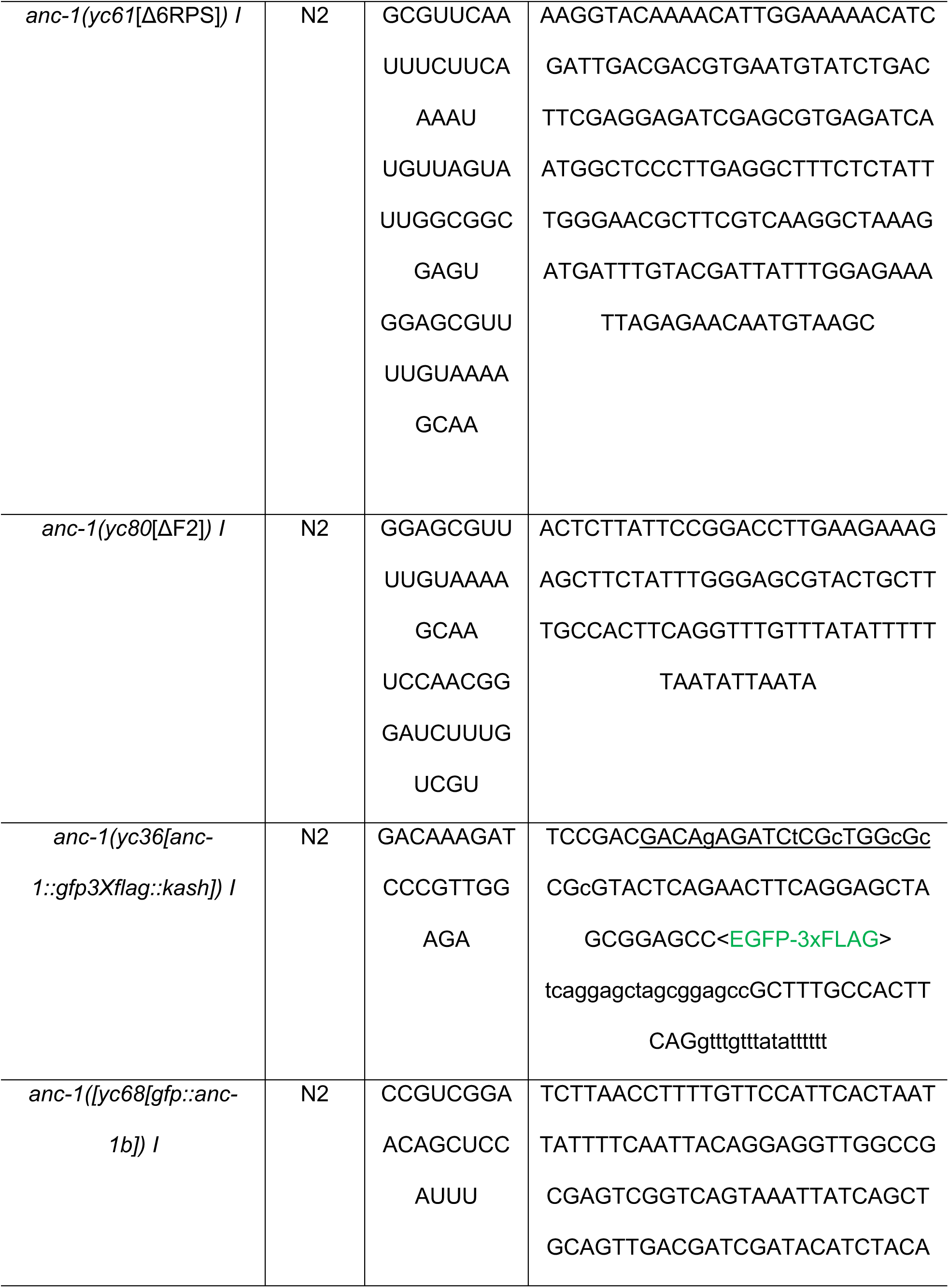
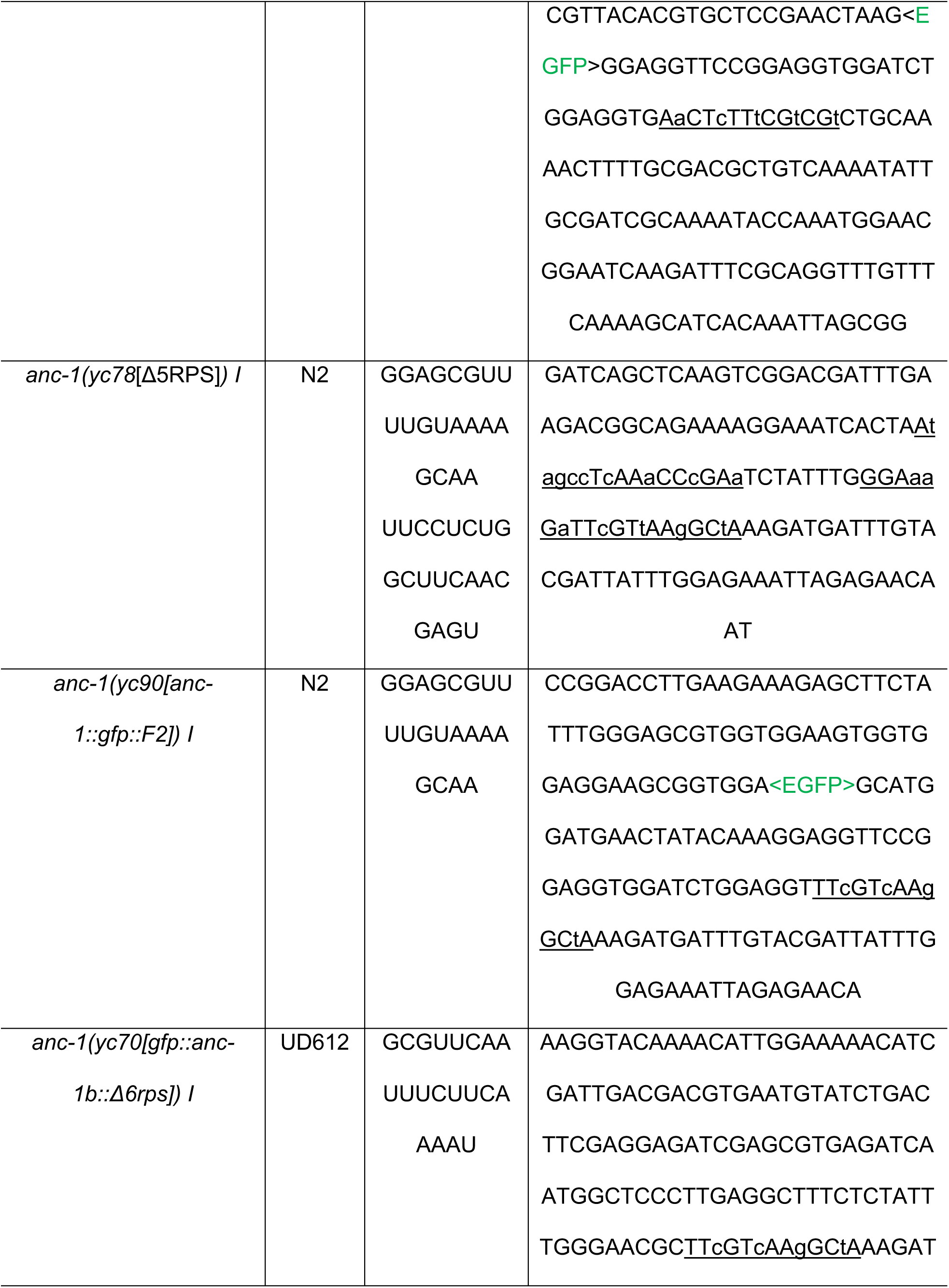
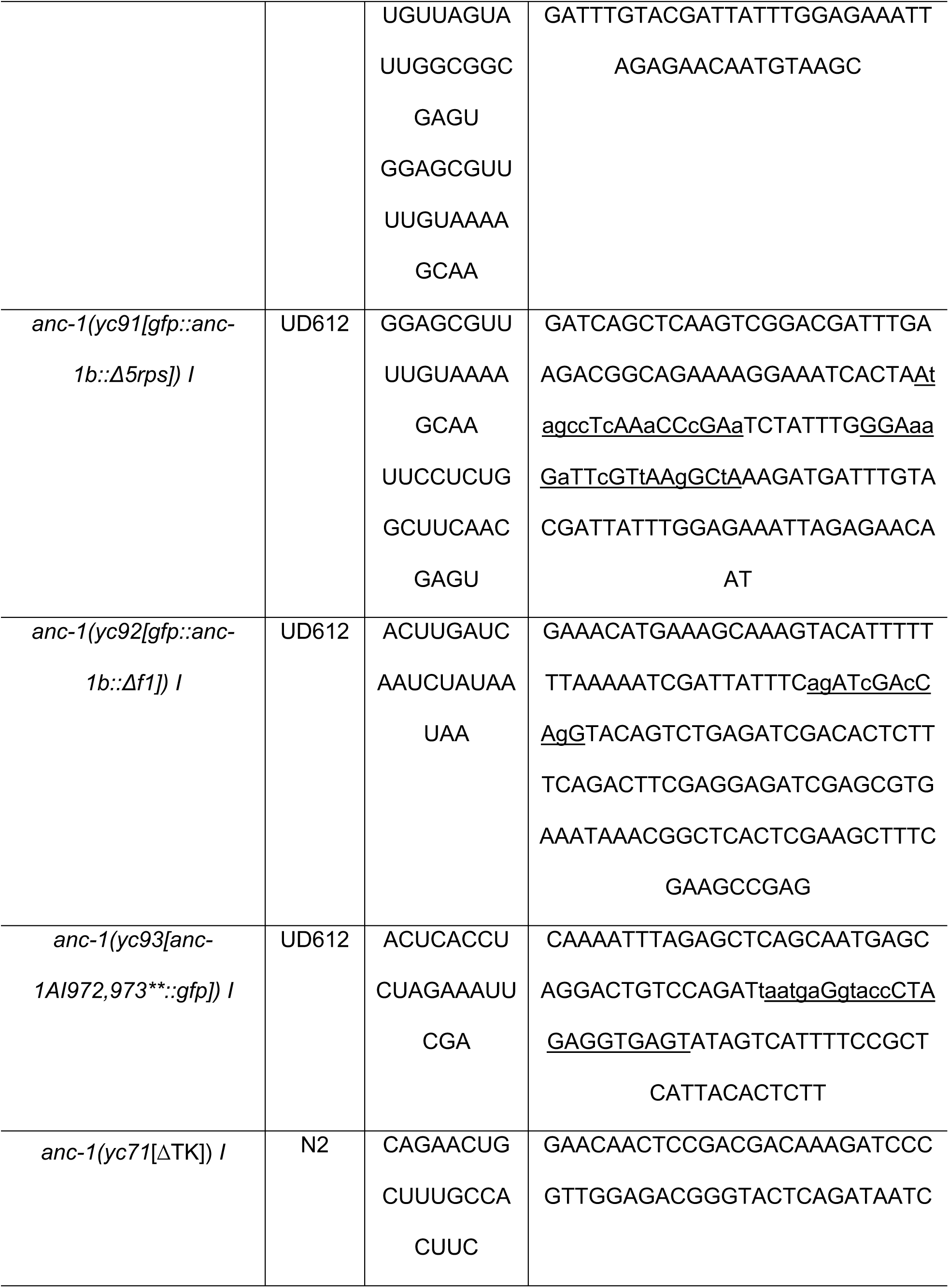
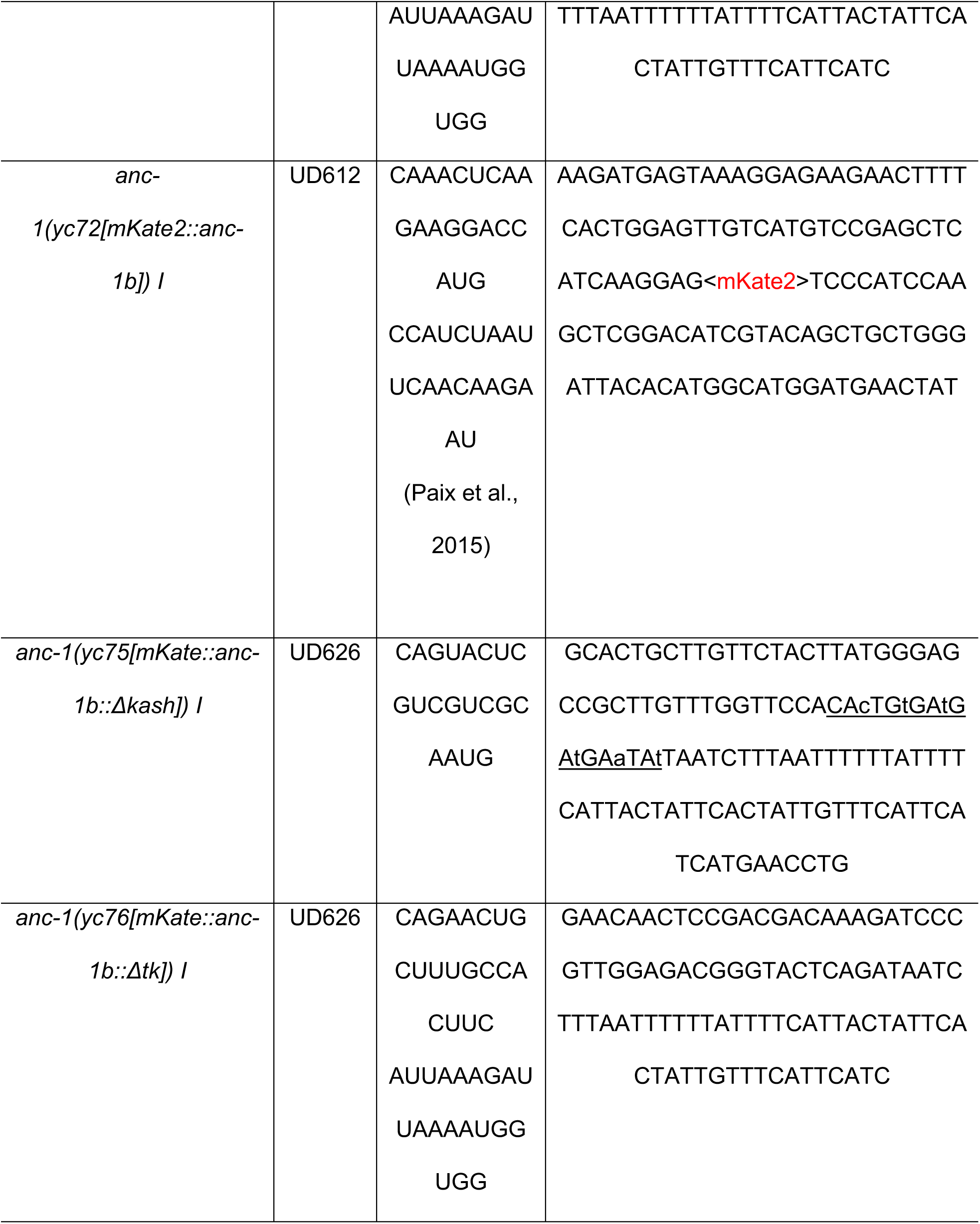
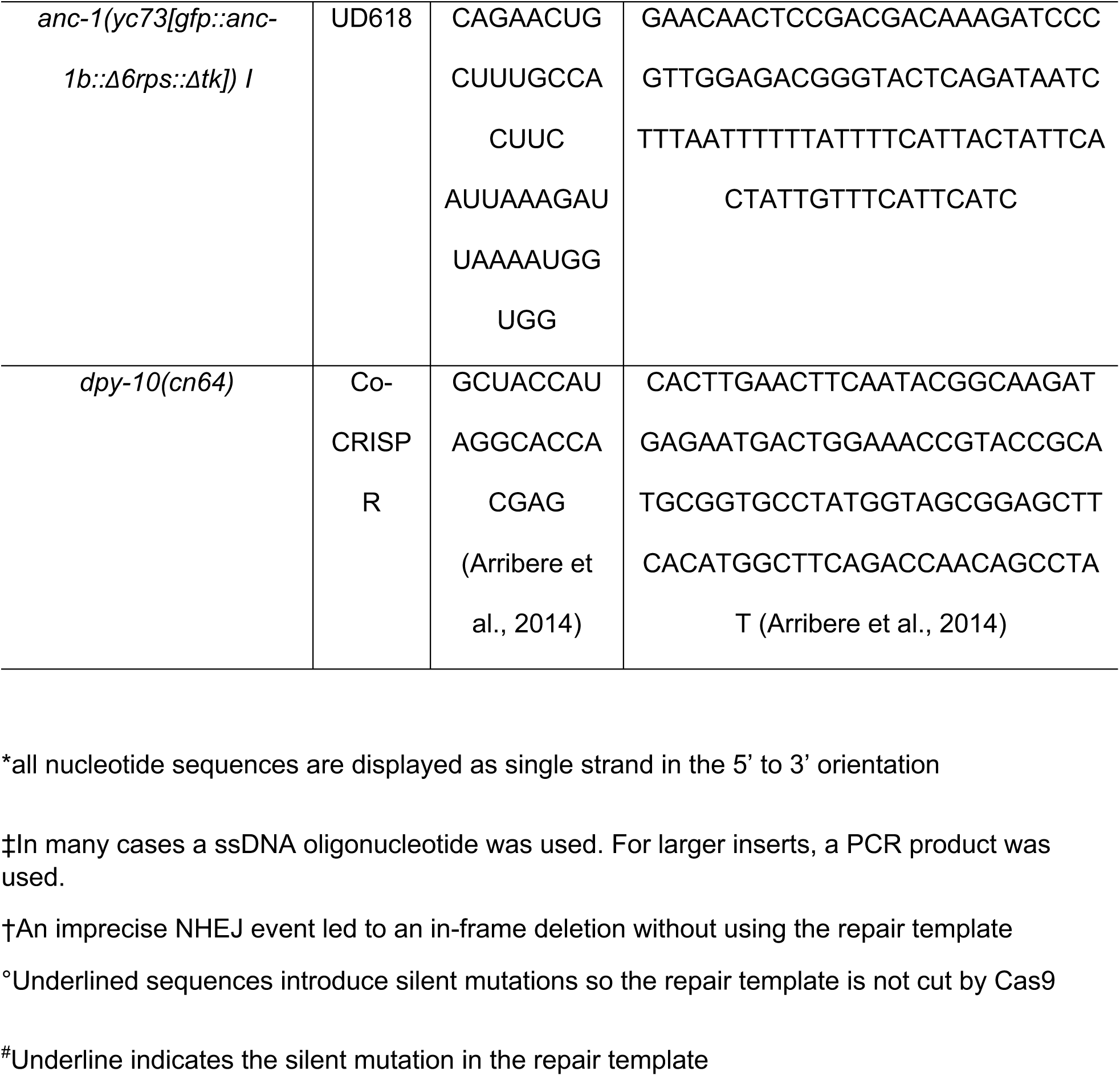
crRNA and repair templates used in this study.

Strain *anc-1[yc41(anc-1::gfp3Xflg::kash,Δch)]I* and *anc-1[yc36(anc-1::gfp3Xflag::kash)]I* were generated by Dickinson Self-Excising Drug Selection Cassette (SEC) method (Dickinson, Pani, Heppert, Higgins, & Goldstein, 2015). In *anc-1[yc41(anc-1::gfp3Xflg::kash,Δch)]I*, both CH domains were deleted (starting with 23KAQK26 and ending with 322QFVR325) and replaced with GFP flanked with 9-residue long linkers (GASGASGAS).

### Microscopy and imaging analysis

Images of the nuclear anchorage, worm body size measurements and *anc-1b* promoter reporter assays were collected with a wide-field epifluorescent Leica DM6000 microscope with a 63× Plan Apo 1.40 NA objective, a Leica DC350 FX camera, and Leica LAS AF software. ANC-1 subcellular localization, the ER marker, and nuclear shape images were taken with a spinning disc confocal microscope (Intelligent Imaging Innovations) with a CSU-X1 scan head (Yokogawa), a Cascade QuantEM 512SC camera (Photometrics), a 100× NA 1.46 objective (Zeiss) and SlideBook software (Intelligent Imaging Innovations). The contrast and levels of the images were uniformly adjusted using ImageJ (National Institutes of Health). Live GFP::KDEL images were acquired at 200 ms intervals using the above spinning-disc confocal system. To quantify ER positioning defects, images from at least ten young adults of each strain were scored blindly by three people. In addition, the “Manual Tracking” plug-in for ImageJ (https://imagej.nih.gov/ij/plugins/track/track.html) was used to track the positions of multiple ER fragments through a time-lapse series. The relative movements between three different spots per animal were measured over time.

For some adult animals, Image J “Stitching” Plug-in were used to stitch images with overlap (Preibisch, Saalfeld, & Tomancak, 2009). The length of L2 larvae, width and length of the adult animals, as well as the circularity and solidity of the nuclei were measured with Image J.

### Statistical evaluation

The nuclear anchorage quantifying data were displayed as scatter plots with means and 95% CI as error bars. Sample sizes were indicated in the figures. Unpaired student t-tests were performed on the indicated comparisons for the nuclear anchorage assay, and Fisher’s exact test was used to determine the difference of the ER anchorage defects between strains. p value less than 0.01 was defined as significant.

## Acknowledgements

We thank Gant Luxton, Erin Cram, Charlotte Kelley, and members of the Starr lab, for helpful discussions and editing of the paper. We thank Barth Grant for sharing an unpublished strain, and Joshua Morgan for suggestions on nuclear shape measurements. We thank Michael Paddy at the MCB Light Microscopy Imaging Facility, which is a UC Davis Campus Core Research Facility, for microscopy assistance. The 3i Marianas spinning disk confocal used in this study was purchased using NIH Shared Instrumentation Grant 1S10RR024543-01. These studies were supported by the National Institutes of Health grants R01GM073874 and R35GM134859 to D.A.S.

---

**Figure S1.**
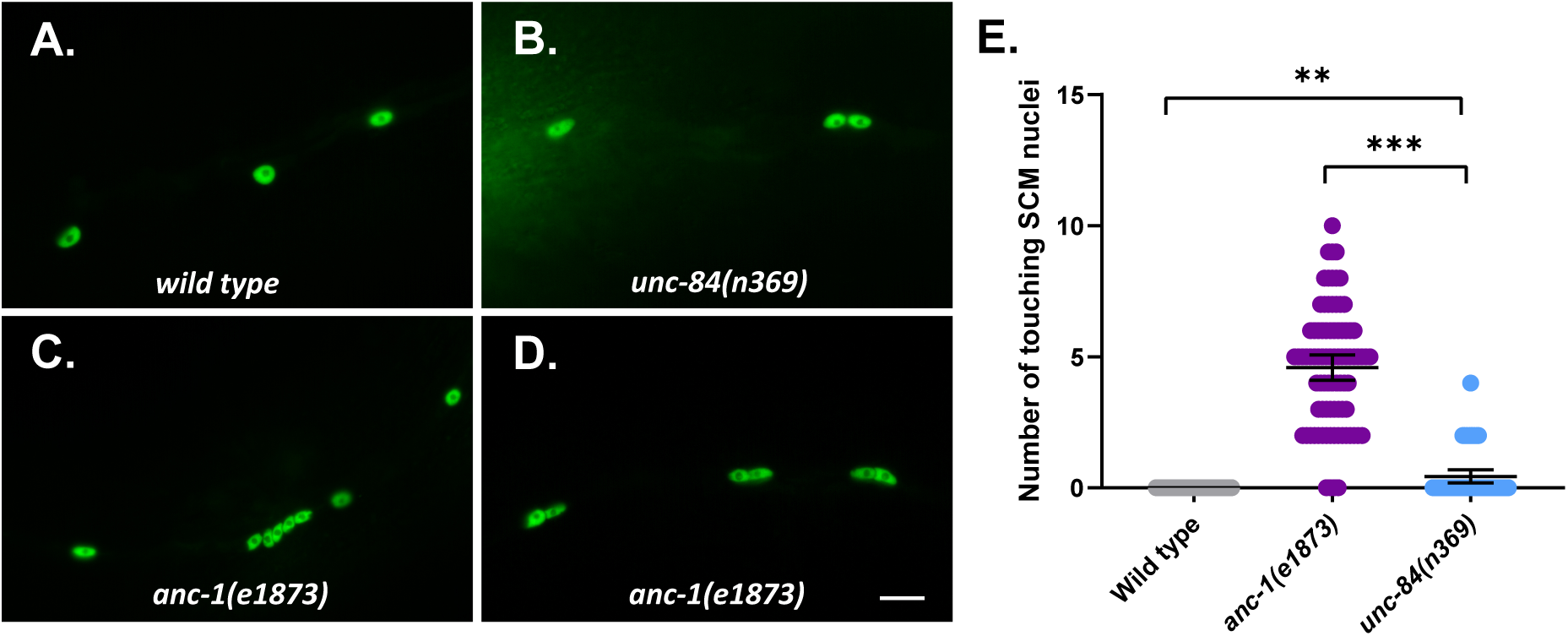
Nuclear anchorage defects in seam cell syncytia. (A-D) Lateral views of *C. elegans* wild type (A), *unc-84(n369)* (B), and *anc-1(e1873)* (C-D) mutants expressing *wIs54[scm::gfp]*. (E) Quantification of the number of touching seam cell nuclei. Each point represents the number of touching nuclei in the seam cell of a young adult animal. Means with 95% CI error bars are shown. Unpaired student two-tail t-test was used for statistical analysis. ns, not significant (p > 0.05); **, P ≤ 0.005. ***, P ≤ 0.001. n ≥ 42 for each strain. Scale bar is 10 µm.

**Figure S2.**
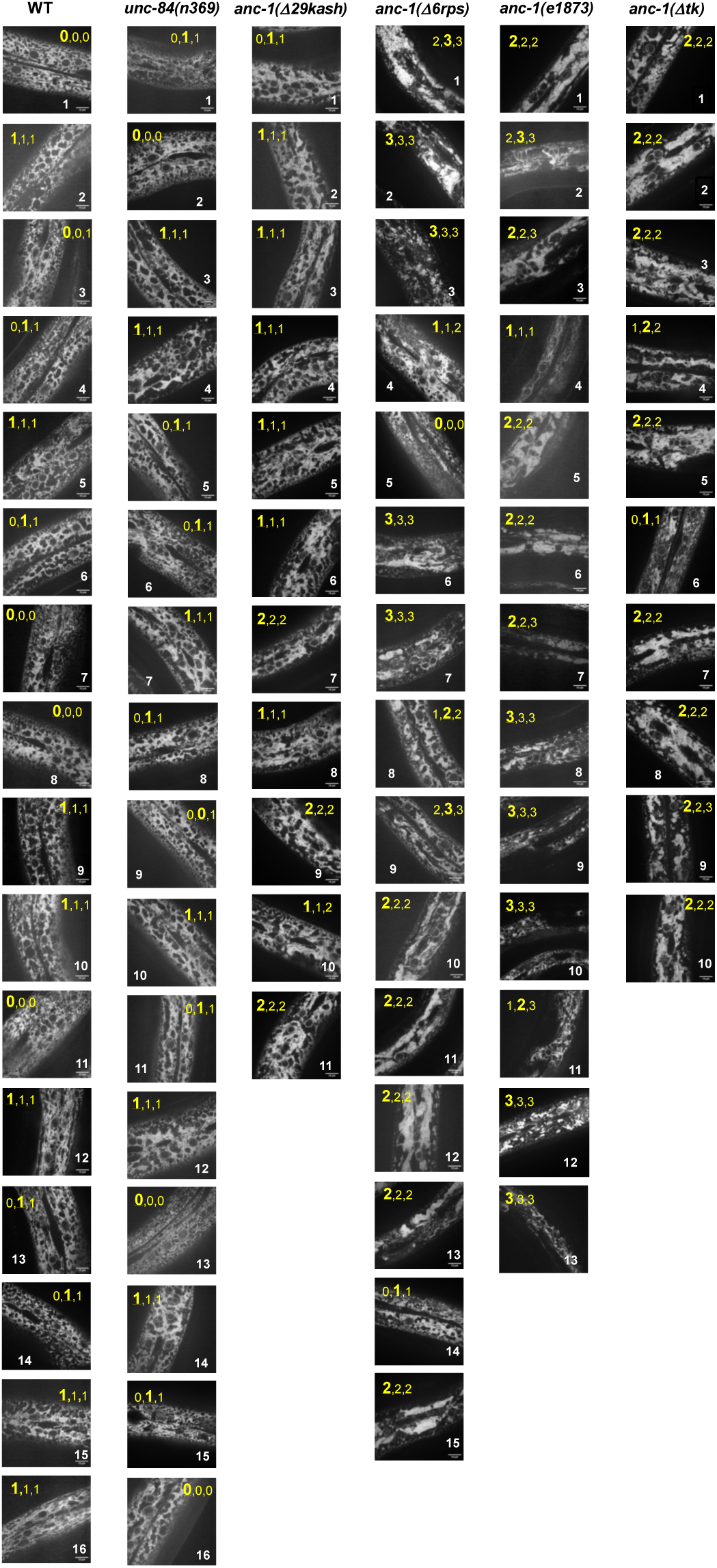
Raw data for ER morphology assays. The ER in hyp7 was labeled with a GFP::KDEL marker. Young adult animals are shown. Each column shows all the images scored from a single genetic strain (labeled on the top). The three numbers in yellow at the top of each image are the scores of three independent investigators giving them bind scores. 0, 1, 2, 3 represents normal, mild, strong and severe ER anchorage defects, respectively. The score given by more than two people was chosen as the final sore and is in a larger font. Scale bars are 10 µm. All the images are at the same scale.

**Movie 1. The ER is anchored in wild-type hyp7**. An example video of the hyp7 ER in young adult wild-type *C. elegans* expressing *pwSi83[p*_*hyp7*_*gfp::kdel]*. Images were captured at the interval of 0.2 s for 10 seconds. Scale bar is 10 µm.

**Movie 2. The ER is unanchored in *anc-1(e1873)* mutant hyp7**. An example video of the hyp7 ER in the young adult *anc-1(e1873)* mutant *C. elegans* expressing *pwSi83[p*_*hyp7*_*gfp::kdel]*. Images were captured at the interval of 0.2 s for 10 seconds. Scale bar is 10 µm.

**Movie 3. ER positioning in *unc-84(null)* mutant hyp7**. An example of one of the most severe ER positioning defects observed is shown in a video of the hyp7 ER in the young adult *unc-84(n369)* mutant *C. elegans* expressing *pwSi83[p*_*hyp7*_*gfp::kdel]*. Images were captured at the interval of 0.2 s for 4 seconds. Scale bar is 10 µm.

